# Immune responses to infection modulate peripheral sympathetic neuron functions

**DOI:** 10.64898/2026.06.25.734445

**Authors:** Pedro Trevizan-Baú, Mitchell T. Ringuet, Maria Daglas, Sapna Devi, Sarah C. Monard, Joon Keit Loi, Ziwei Luo, Alicia Weier, Lachlan Dryburgh, Yannick O. Alexandre, Hyun Jae Lee, Shihan Li, Daniel Rawlinson, Emily A. Canner, Dominik Schienstock, Harry L Horsnell, Thomas N. Burn, Zoe Fransos, Sam Mohammed, Lukasz Kedzierski, Isabelle J. H. Foo, Madeleine R. Di Natale, Marcela de Lima Moreira, Juan C. Molero, Garron T. Dodd, Katherine Kedzierska, Laura K. Mackay, Ashraful Haque, Erica K. Sloan, Jan Schröder, Robin M. McAllen, John B. Furness, Scott N. Mueller

## Abstract

The central nervous system interprets inflammatory signals in the body and directs the modulation of inflammatory responses by reflexively engaging peripheral sympathetic neurons^1,2^. This includes sympathetic neurons that innervate the spleen, which can regulate immune functions^3–5^ and modulate inflammation^6–8^. Yet, it is unclear if neuroimmune interactions involve specialised immunoregulatory sympathetic neurons, and if the immune system can reciprocally regulate peripheral sympathetic neurons to control these responses. Using retrograde tracing and single-cell transcriptomics, we find that spleen-innervating neurons are heterogeneous but do not exhibit a distinct transcriptional program indicative of specialisation for immune communication. However, we report that immune responses induced by pathogens can regulate postganglionic sympathetic neuron functions. Cytokines produced by immune cells downregulate expression of the neurotrophin nerve growth factor in spleen mesenchymal cells, leading to organ-specific sympathetic nerve retraction from the spleen. Concurrently, splenic type I interferon signalling induces inflammatory gene expression in neurons and suppresses neuron excitability. Chemogenetic activation of sympathetic neurons demonstrates an impaired anti-inflammatory capacity in the spleen during infection. These results reveal regulation of sympathetic neuronal functions by the immune system, which could support optimal generation of immune responses against pathogens.

## Introduction

The immune system and nervous system both underpin body physiology and play important roles in protection against, and recovery from, infections. Subtypes of sensory neurons have been observed to engage in bi-directional crosstalk with immune cells in various tissues and settings that influence the outcomes of immune responses^9,10^. This includes sensory neurons that can support immune responses in the spleen^11,12^. Conversely, studies have shown that sympathetic immunomodulatory neurons control inflammation, including the participation of defined brain regions in activating peripheral neuroimmune circuits^1,13,14^ and a role for sympathetic nerves in the spleen in endogenous inflammatory reflexes^6,15^. The influence of sympathetic neuron-derived signals on immune cells is becoming better defined^8,16,17^, however the capacity for feedback regulation of peripheral sympathetic neurons by the immune system is largely unknown.

Sympathetic neurons can be divided into different functional classes based on the capacity to modulate tissue physiology, for example vasoconstrictor neurons, pupillodilator neurons and cardio-accelerator neurons^18,19^. It has been proposed that immunomodulatory sympathetic neurons, including neurons that innervate the spleen, may be functionally distinguishable and represent a sub-class that is specialised for neuroimmune communication^20,21^. Yet, these lymphoid tissue projecting neurons have not been isolated. Moreover, knowledge of the phenotypic and transcriptional diversity of peripheral sympathetic neurons remains limited. Here we characterised heterogeneous spleen-innervating sympathetic neurons and identify a regulatory pathway that is activated during infection via the spleen to control sympathetic neuron functions. Our results provide new insights into the mechanisms by which peripheral sympathetic neurons respond to infection and suggest an additional control step in the inflammatory reflex that enables the immune system to modulate sympathetic neurons to help orchestrate immunity.

## Results

### Characterisation of spleen-innervating sympathetic neurons

The axons of sympathetic post-ganglionic neurons, that were identified through their immunoreactivity for tyrosine hydroxylase (TH), enter the spleen at the hilum and follow the splenic artery as it branches and traverses the white pulp. TH-containing nerve fibres were imaged in thick sections of mouse spleen and observed in a dense meshwork around central arteries (CD31^+^ vessels, white) in the T cell zone (Fig. 1a). Nerve branches extended from the peri-arterial plexuses through T cell zones and some reached B cell follicles. As anticipated^22^, TH^+^ axons were in close proximity to lymphocytes (Fig. 1b(i)). Unexpectedly, examination of the stromal niche in the white pulp using CCL19-tdTomato transgenic mice that contain fluorescently labelled fibroblastic reticular cells revealed that nerve branches consistently follow the stromal cell networks in the T cell zone (Fig. 1b(ii)). This anatomical arrangement indicates ample potential for functional interactions between spleen-innervating sympathetic neurons and both stromal and immune cell networks. Therefore, we conducted experiments to establish the molecular identity of spleen sympathetic neurons to determine if transcriptionally distinct sympathetic neuron populations innervate the spleen, and to predict potential mechanisms by which they modulate splenic functions.

**Figure 1:**
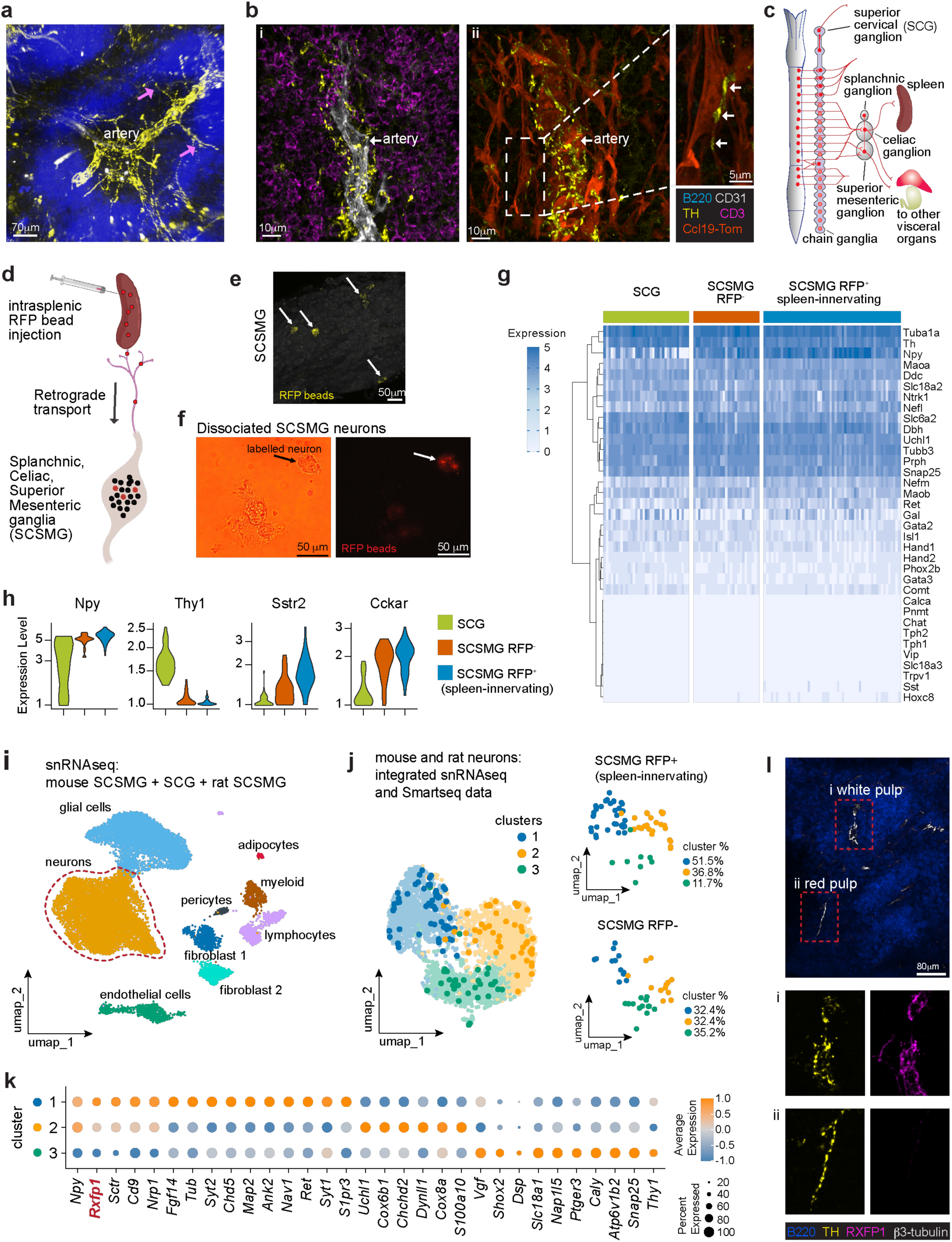
Heterogeneous sympathetic neurons innervate the spleen but do not specialise for neuroimmune communication. **a**) Vibratome section of mouse spleen stained for TH (yellow; nerve fibers), B220 (blue, B cells) and arteries (CD31, white). TH^+^ axons branch from the peri-arterial plexus throughout the white pulp (arrows). **b**) Spleen white pulp showing TH^+^ axons in association with a CD31^+^ central artery and (i) T cells (CD3ε, purple), and (ii) with Ccl19-tdTomato^+^ stromal cells (red). **c**) Schematic of paravertebral and prevertebral sympathetic ganglia highlighting the SCSMG complex and the SCG. **d**) Schematic of approach used for spleen neuron tracing. RFP microspheres were injected into spleens and SCSMG isolated from mice 7-12 days later. **e**) Imaging of isolated SCSMG showing RFP bead-labelled neurons (arrows). **f**) Imaging of dissociated SCSMG neurons depicting cell morphology (brightfield, left) and the presence of RFP beads in a labelled neuron (fluorescence, right). **g**) Heatmap of selected neuronal marker genes amongst 35 SCG, 27 unlabelled (RFP^-^) SCSMG and 56 labelled (RFP^+^) SCSMG neurons from mice profiled by scRNA-seq. **h**) Expression of selected genes in indicated groups of neurons. **i**) Droplet-based snRNA-seq of SCSMG from mice and rats and from published mouse SCG^25^, representing 16,310 cells. **j**) UMAP projections showing 3353 neurons profiled by snRNA-seq and 140 neurons profiled by scRNA-seq that were integrated together. Neurons profiled by scRNA-seq are highlighted on the plots, overlaid onto the snRNA-seq data and shown separately on the right to distinguish the unlabelled (RFP^-^) and labelled (RFP^+^) SCSMG neurons. The proportion of cells in each sample group that were assigned to clusters is shown. **k**) Differentially expressed genes amongst the three neuron clusters. **l**) Confocal microscopy of spleen sections from Rxfp1-tdTomato mice showing innervation in the (i) white pulp and (ii) red pulp.

The somas (cell bodies) of mouse spleen-innervating sympathetic neurons were located in prevertebral sympathetic ganglia (Fig. 1c) by back-labelling using intrasplenic injection of red fluorescent protein (RFP)-labelled microspheres (Fig. 1d). The microspheres were retrogradely transported to neuron somas and visualised in the mouse sympathetic splanchnic-celiac-superior mesenteric ganglion (SCSMG) complex (Fig. 1e)^23^. As expected^24^, retrogradely labelled cell bodies were not located in the superior cervical ganglia (SCG), which are paravertebral sympathetic ganglia that innervate targets in the head and neck. Enzymatic dissociation of the isolated SCSMG and visualisation of fluorescently labelled neurons was followed by manual isolation of single labelled (RFP^+^, spleen-innervating) and unlabelled (RFP^-^) neurons from the SCSMG as well as control neurons from the SCG (Fig. 1f). We performed scRNA-seq using the Smart-seq2 protocol on 56 RFP^+^ spleen-innervating neurons, 27 control RFP^-^ SCSMG neurons and 35 RFP^-^ SCG neurons isolated using this approach.

The profiled neurons expressed canonical genes involved in nervous system development, ion channels, G-protein coupled receptors and synapses (Extended Data Fig. 1a), including catecholamine biosynthesis pathway genes (*Th*, *Ddc*, *Dbh*), the neurotransmitter transporters Nat1 (noradrenaline transporter 1, *Slc6a2*) and Vmat2 (vesicular monoamine transporter 2, *Slc18a2*), as well as genes expressed in peripheral neurons, peripherin (*Prph*), β3-tubulin (*Tubb3*) and synaptosome associated protein 25 (*Snap25*) (Fig. 1g). The sympathetic neurons expressed the monoamine oxidases *Maoa* and *Maob*, and catechol-O-methyltransferase (*Comt*) that are required for the metabolism of catecholamines. Importantly, genes defining visceral sensory neurons (*Calca*, *Trpv1*), cholinergic neurons (*Chat*, *Slc18a3*) or serotonergic neurons (*Tph1*, *Tph2*) were not expressed, confirming that all neurons that we profiled were noradrenergic. We also performed retrograde tracing and scRNAseq on SCSMG neurons in the rat, a species frequently used to study the nervous system. This revealed that sympathetic neurons in SCSMG are transcriptionally conserved between rats and mice (Extended Data Fig. 1b).

Comparison of mouse SCSMG and SCG neurons identified expression of neuropeptide Y (*Npy*) in all spleen-innervating neurons, whereas 40% of SCG neurons expressed low or negligible *Npy* (Fig. 1g and h). Furthermore, SCSMG neurons differentially expressed the homeobox genes *Hoxc8* and *Hoxc9*, the angiotensin receptor *Agtr1a*, the oxygen binding protein neuroglobin (*Ngb*), somatostatin receptor 2 (*Sstr2*) and the cholecystokinin A receptor (*Cckar*) (Fig. 1h and Extended Data Fig. 2a). Conversely, SCG neurons expressed the transcription factors *Rorc* and *Hoxc4*, as well as the glycoproteins *Thy1* and *Scube1*. This signifies specialisation of the transcriptome of noradrenergic neurons located in the SCSMG and SCG. However, comparison of RFP^+^ and RFP^-^ neurons from SCSMG revealed largely comparable gene expression, suggesting similar transcriptional diversity (Extended Data Fig. 2b).

### Spleen-innervating neurons are heterogeneous but not specialised

To further interrogate spleen-innervating neurons we performed single nucleus RNA sequencing (snRNAseq) on SCSMG from mice and rats and incorporated published snRNA-seq data from mouse SCG^25^ (Fig. 1i). All cell type clusters were present in mouse SCSMG and SCG as well as the rat SCSMG tissues (Extended Data Fig. 2c). Neurons were defined by expression of genes including *Npy*, *Tubb3*, *Th* and *Prph*, and glial cells by canonical genes including *Scn7a*, *Sox10* and *S100b* (Extended Data Fig. 2d). We annotated two fibroblast clusters (*Dcn*, *Lum*, *Fap*), that included a *Pdgfra* and *Col15a1* cluster and a *Itga6* and *Thbs4* expressing cluster. We also identified endothelial cells (*Flt1, Tek, Pecam1*), pericytes (*Myh11, Rgs5*), adipocytes (*Lpl*, *Pparg*) and immune cell clusters that included myeloid cells (*Lyz2, Fcer1g*) and lymphocytes (*Lck, Pax5*). Next, we sub-clustered neurons and integrated these with our Smartseq profiled neurons, which revealed three neuronal clusters that were present in all samples irrespective of species, ganglion or isolation protocol (Fig. 1j and Extended Data Fig. 2e).

Spleen-innervating neurons were enriched in neuron clusters 1 and 2 and were characterised by higher expression of genes that included *Npy* and relaxin family peptide receptor 1 (*Rxfp1*), a receptor for the hormone relaxin (Fig. 1k). A smaller proportion of spleen neurons in cluster 3 expressed the transcription factor *Shox2* and the protein *Bmp3*. GO term analysis revealed enrichment for pathways involved in axonogenesis and nervous system development by neurons in cluster 2, whereas neurons in the adjacent cluster 1 were enriched for aerobic respiration and microtubule genes (Extended Data Fig. 2f). We further validated spleen-innervating neuron clusters by integrating our Smartseq profiled neurons with recently published scRNAseq analysis of SCSMG^26^. This analysis also identified 3 neuron clusters, including 2 *Rxfp1^+^*clusters that were enriched in spleen-innervating neurons as well as a smaller *Shox2^+^* cluster (Extended Data Fig. 2g, h). Together, these data identify transcriptionally heterogenous sympathetic neurons innervating the spleen.

A recent study identified organ-specific sympathetic innervation of visceral organs by either RXFP1^+^ or SHOX2^+^ SCSMG neuronal subtypes^27^. Since our retrogradely traced spleen neurons included both neuronal subtypes, we validated RXFP1 expression using reporter mice. This revealed RXFP1 expression by white pulp sympathetic nerves, whereas a subgroup of red pulp nerves were RXFP1^-^ (Fig. 1l). These results demonstrate that the sympathetic neurons that innervate the spleen are heterogeneous, possibly indicating distinct target cells or functions. But these neurons are not molecularly distinct from sympathetic neurons populations that innervate other visceral organs, indicating that the spleen is not innervated by neurons with a discrete immunoregulatory transcriptional profile.

### Retraction of splenic sympathetic innervation during infections

Sympathetic nerves are reflexively activated during immune challenge and can regulate immune responses in the spleen^6–8^. Since our experiments did not identify transcriptionally unique immunoregulatory neurons innervating the spleen, we postulated that other mechanisms might exist to enable context-dependent regulation of neuroimmune interactions during immune responses. To investigate this, we infected mice with lymphocytic choriomeningitis virus (LCMV) and examined axons in the spleen after infection (Fig. 2a). We performed tissue clearing using a modified Shanel method and immunofluorescence staining of whole-mount spleens to visualise sympathetic nerves in 3D. Uninfected mice had dense arborisation of TH^+^ innervation throughout the spleen (Fig. 2b, c). Unexpectedly, LCMV infection induced a substantial retraction of sympathetic innervation in the spleen, marked by almost complete axon loss in the white pulp and red pulp, including the dense innervation around intrasplenic blood vessels (Fig. 2c). Spleen sympathetic nerves retracted to the splenic artery in the hilum where TH^+^ staining persisted after infection (Fig. 2b and Extended Data Fig. 3a). We quantified splenic sympathetic innervation by staining sections with the nerve markers TH, synaptophysin and β3-tubulin in conjunction with CD31 and B220 to identify blood vessels and B cell follicles, respectively (Fig. 2d). This revealed that reduction in splenic sympathetic innervation began by day 3 of infection, with almost complete loss of innervation by day 8, corresponding with the peak of the immune response (Fig. 2e). Restoration of splenic innervation after LCMV infection occurred over several weeks. To confirm that our observations reflected loss of nerve fibres, not just TH, from the spleen, we generated TH-cre/Ai14 mice with TdTomato labelled adrenergic neurons. This confirmed that infection resulted in a significant loss of sympathetic innervation from the spleen (Extended Data Fig. 3b).

**Figure 2:**
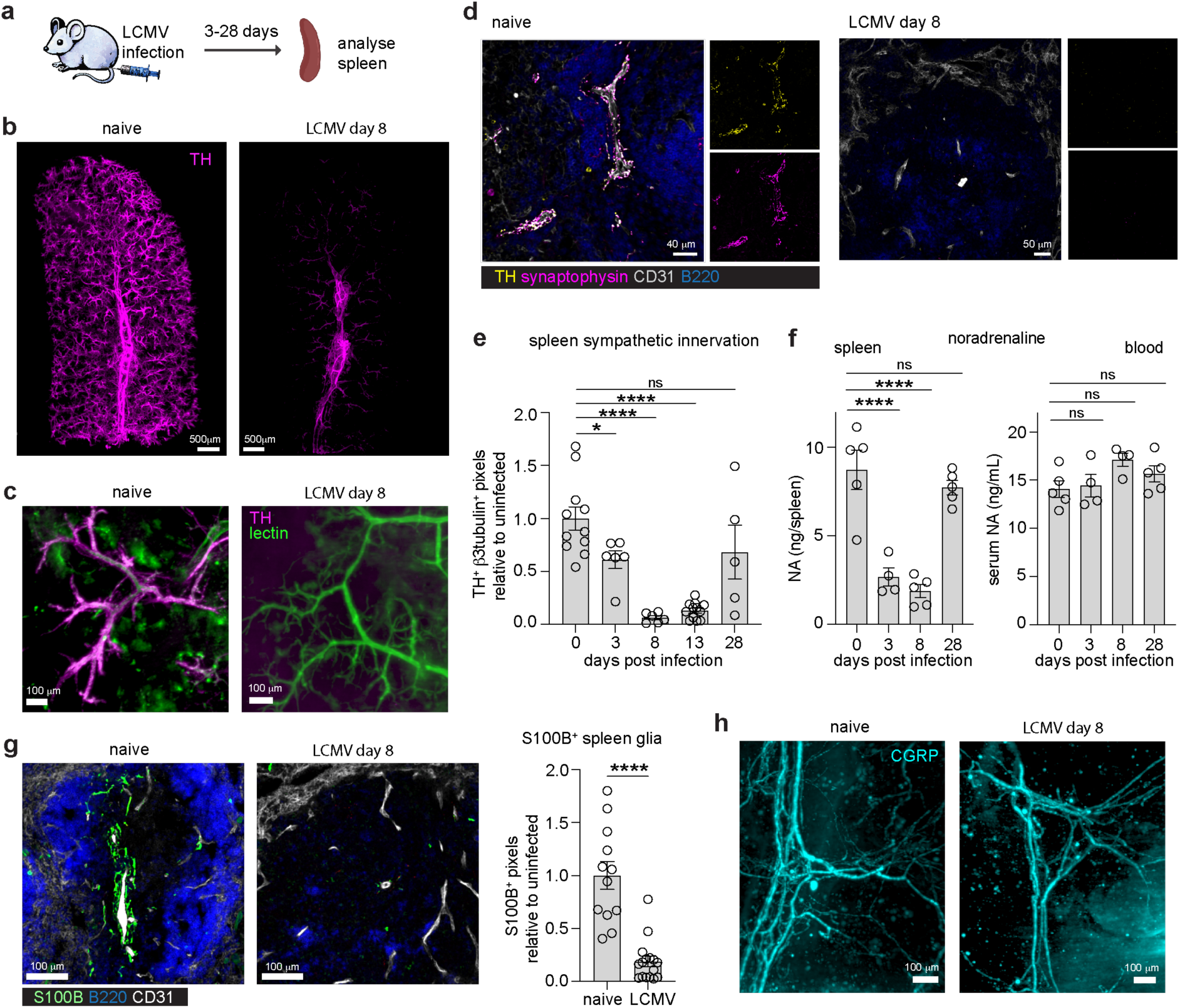
Infections induce sympathetic neuropathy of the spleen. **a**) Mice were infected i.p. with LCMV and spleens analysed between 3 and 28 days later. **b**) 3D view of TH staining in cleared whole-mount spleens from naïve and day 8 LCMV mice. **c**) Optical sections of tomato-lectin labelled blood vessels and TH staining in cleared spleens from naïve and day 8 LCMV mice. **d**) Confocal imaging of spleen sections from naïve and day 8 LCMV mice, stained for TH, synaptophysin, CD31^+^ endothelial cells and B220^+^ B cells. **e**) Quantitation of sympathetic innervation in spleen sections from naïve and LCMV infected mice. n = 5-12 mice per group. **f**) Measurement of noradrenaline (NA) in spleen tissues and blood (serum) from naïve mice (day 0) and 3, 8 and 28 days after LCMV infection. n = 4-5 mice per group. **g**) Staining for S100B^+^ glial cells, CD31 and B220 in spleen sections (images) and quantitation of glial cell density in naïve and LCMV infected mice (graph). n = 12-16, 2 regions from 6-8 mice per group. **h**) Optical sections from light sheet imaging of CGRP staining in cleared whole-mount spleens from naïve and day 8 LCMV mice. Graphs show representative data (mean ± SEM) from one of two independent experiments (f) or two independent experiments (e, g). *P < 0.05, ****P < 0.0001, ns, non-significant; by Brown-Forsythe & Welch ANOVA (e) one-way ANOVA (f) or unpaired t-test (g).

Visualisation of catecholamines in spleen sections using sodium-potassium-glyoxylic acid (SPG) histofluorescence identified expected dense staining around intrasplenic arteries in uninfected mice (Extended Data Fig. 3c). At the peak of the immune response in the spleen 8 days after LCMV administration, catecholamines were largely undetectable within the spleen. Measurement of splenic noradrenaline (NA) levels confirmed the substantial reduction in this sympathetic neurotransmitter after infection (Fig. 2f). Reduction in spleen NA occurred as early as 3 days after infection and returned to approximately normal levels within one month, but this loss was not associated with changes in circulating NA measured in the blood (Fig. 2f). Furthermore, sympathetic neuropathy was not detected in the myenteric plexus in the gastrointestinal tract, another site of LCMV infection that is also innervated by neurons located in the SCSMG, since TH^+^ innervation of the myenteric plexus was unchanged (Extended Data Fig. 3d).

We recently characterised a subset of glial cells that ensheath sympathetic nerves in the spleen^28^. Staining for S100B^+^ glial cells revealed that the spleen glial cell network was significantly reduced following LCMV infection, indicating loss concomitant with axon retraction (Fig. 2g). Recent studies have demonstrated that sensory neurons innervate spleen blood vessels, particularly in the hilum and the vessels branching from the hilum artery^11^. The neuropeptide calcitonin gene-related peptide (CGRP) is produced by spleen nociceptors and can enhance B and T cell responses in the spleen^11,12^. To determine if sensory nerves also retracted from the spleen after infection, we imaged cleared whole-mount spleens that were stained for CGRP. This identified intact sensory nerve fibres in infected mice that remained concentrated in the hilum (Fig. 2h and Extended Data Fig. 3e). Thus, we identified a pronounced organ-specific sympathetic neuropathy in the spleen during virus infection. We also examined TH^+^ splenic innervation in mice infected with the bacterial pathogen *Listeria monocytogenes*, and the malarial parasite *Plasmodium berghei* ANKA. Both infections induced a significant reduction in spleen sympathetic innervation (Extended Data Fig. 3f).

### An immune-stromal circuit modulates splenic innervation

Because changes to spleen innervation paralleled the immune response to infection, we hypothesised that axon loss was driven by components of the immune response. Antibody-mediated depletion of CD8^+^ T cells did not significantly alter the loss of sympathetic innervation in the spleen after infection (Fig. 3a). By contrast, we observed retention of approximately 25% of innervation when CD4^+^ T cells or both CD8^+^ and CD4^+^ T cells were depleted. Almost 50% of spleen TH^+^ innervation was retained following the depletion of NK cells (Fig. 3b). In contrast, depletion of macrophages did not impact loss of spleen innervation (Extended Data Fig. 4a). This suggested contributions by several cell types that are involved in anti-viral immunity, and possible additional contributions by other immune cells. Investigation of potential inflammatory mediators of nerve loss found no role for type I interferon signals (Fig. 3c), however infection of TNF or IFNγ knockout mice, or TNF/IFNγ double knockout mice, revealed necessary expression of both TNF and IFNγ for the retraction of spleen innervation (Fig. 3d,e). Treatment with anti-TNF and anti-IFNγ antibodies throughout infection confirmed that these cytokines were responsible for loss of spleen innervation (Extended Data Fig. 4b).

**Figure 3:**
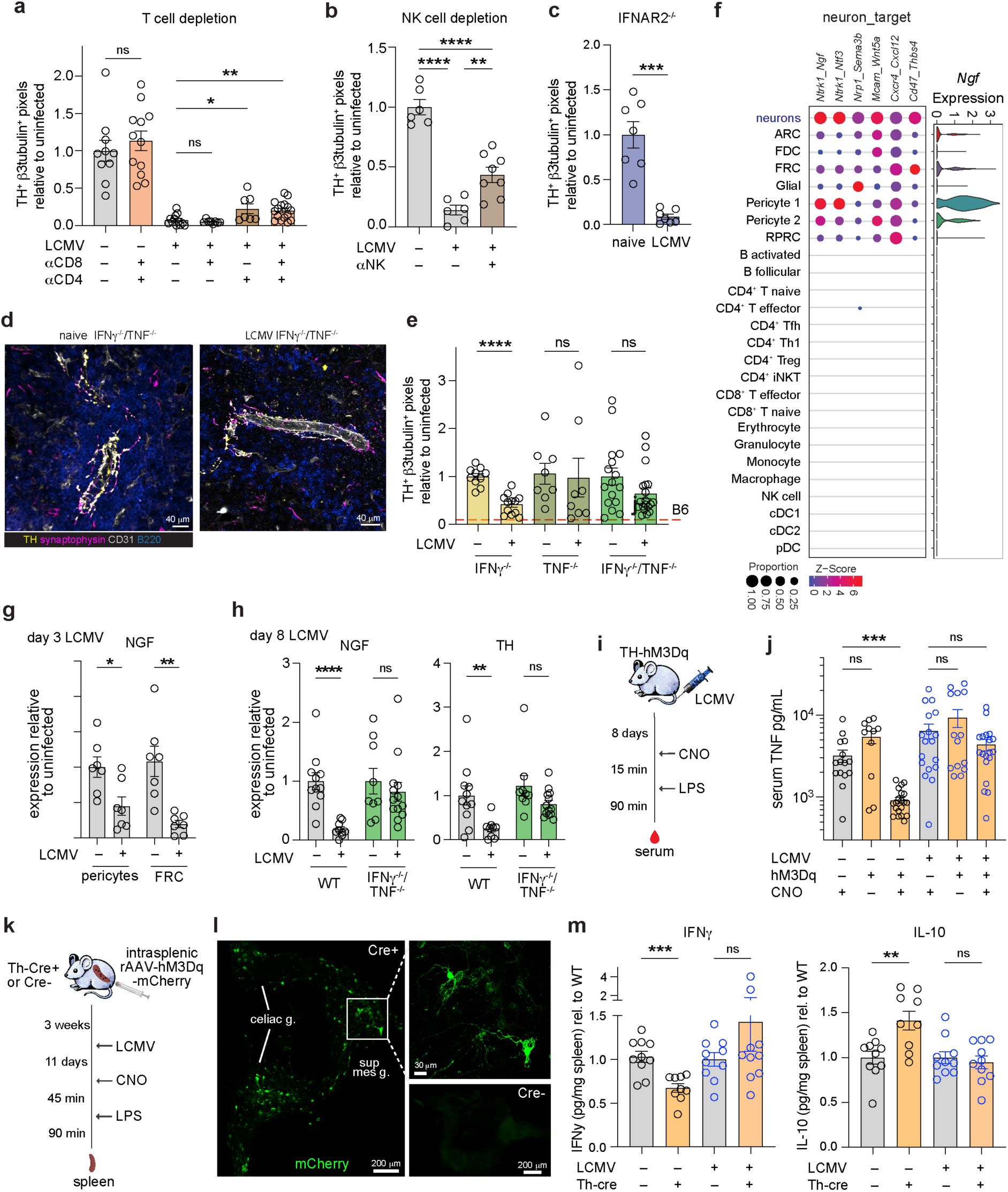
An immune-stromal-nerve circuit modulates sympathetic immunoregulation in the spleen. **a, b**) Depletion of immune cells and assessment of spleen innervation. Quantitation of sympathetic innervation in spleen sections from naïve and LCMV infected mice treated with CD4 or CD8 T cell depleting antibodies (**a**) or anti-NK1.1 antibodies (**b)** prior to infection. N = 6-15 mice per group. **c**) Assessment of spleen innervation in naïve or day 8 LCMV infected IFNAR2^-/-^ mice. n = 6-7 mice per group. **d**) Spleen sections from naïve and day 8 LCMV IFNγ^-/-^/TNF^-/-^ mice, stained for the indicated markers. **e**) Quantitation of sympathetic innervation in spleen sections from naïve and LCMV infected IFNγ^-/-^, TNF^-/-^ and IFNγ^-/-^/TNF^-/-^ mice. Average level of innervation in WT mice at day 8 is shown by the red dotted line. n = 8-18 mice per group. **f**) Top receptor-ligand interactions predicted between spleen neurons and spleen stromal and immune cells (dotplot) and expression of *Ngf* by the indicated spleen cell types (violin plot). ARC, adventitial reticular cell; FDC, follicular dendritic cell; FRC, fibroblastic reticular cell; RPRC, red pulp reticular cell. **g**) Expression of NGF by spleen pericytes and FRC from naïve and day 3 LCMV mice. **h**) Expression of NGF and TH in spleens from naïve and day 8 LCMV infected WT (B6) and IFNγ^-/-^/TNF^-/-^ mice. n = 9-13 mice per group. **i**) Experimental activation of the sympathetic nervous system to modulate TNF production in response to LPS administration. **j**) Naïve or day 8 LCMV infected TH-Cre.hM3Dq-DREADD mice (Cre^+^) or control mice (Cre^-^) were examined after CNO or PBS treatment and administration of LPS. TNF was measured in serum 90 min after LPS administration. n = 11-20 mice per group. **k**) Intrasplenic injection of rAAV-hM3Dq-mCherry in TH-Cre^+^ or Cre^-^ mice 17d before LCMV infection. On day 11 naïve or infected mice were treated with CNO and then LPS before isolation of spleens 90 min later. **l**) Imaging of mCherry expressing neurons in cleared SCSMG whole-mounts of naive TH-Cre^+^ or Cre^-^ mice. **m**) IFNγ and IL-10 were measured in spleen supernatants 90 min after LPS administration. n = 9-10 mice per group. Graphs show representative data (mean ± SEM) from two independent experiments (a-c, e, g-h, m) or 3-4 independent experiments (j). *P < 0.05, **P < 0.01, ***P < 0.001, and ****P < 0.0001, ns, non-significant; by Brown-Forsythe & Welch ANOVA (a, e, g, j), one-way ANOVA (b, m), Welch’s t-test (c) or Kruskal-Wallis tests (h).

We next investigated potential pathways contributing to the loss of spleen innervation. To characterise the molecular interaction networks that exist between spleen-innervating neurons and the diverse cell types in the spleen, we identified genes encoding ligand and receptor pairs expressed by spleen neurons and immune and stromal cells using our previously published scRNA-seq datasets^28,29^ (Extended Data Fig. 4c,d). We restricted this analysis to genes expressed in at least 50% of cells and ranked the cell types by the number of predicted molecular interactions (Extended Data Fig. 4e). Subsets of mesenchymal stromal cells expressed the highest number of predicted interacting partners with the sympathetic neurons, reflecting our observation of interactions between spleen nerves and fibroblasts (Fig. 1b). Communication between spleen neurons and immune cells was dominated by β2 adrenergic receptor (*Adrb2*) interactions (Extended Data Fig. 4f). Neuron to stromal cell communication included *Npy* and fibroblast growth factor (*Fgf1*) signalling from spleen neurons to *Npy1r* and *Fgfr1* expressed by splenic stromal cells. Prominent signals predicted to be received by spleen neurons from spleen stromal cells include the chemokine *Cxcl12* that binds to *Cxcr4* and could assist in neuron migration and survival^30^, and the growth factors nerve growth factor (*Ngf*) and neurotrophin 3 (*Ntf3*), expressed by pericytes and fibroblastic reticular cells, that bind to TrkA (*Ntrk1*), the high-affinity tyrosine kinase receptor on neurons (Fig. 3f).

It has been shown that mice lacking NGF do not have sympathetic innervation in the spleen, but innervation remains on the splenic artery^31^. NGF provides retrograde trophic support for sympathetic axons in target organs through the TrkA receptor^32^. Our scRNA-seq data indicates that expression of NGF is concentrated in stromal cells that form the perivascular niche and is consistent with the spatial localisation of NGF mRNA described in mouse spleen^33^. We hypothesised that disrupted innervation after infection might be driven by changes in NGF availability. In support of this, we found that expression of NGF was significantly downregulated in pericytes and FRCs in the spleen by day 3 of LCMV infection (Fig. 3g). We tested if TNF and IFNγ contributed to the downregulation of NGF. The absence of TNF and IFNγ was able to rescue NGF expression in the spleen after infection (Fig. 3h), consistent with retention of sympathetic innervation in TNF^-/-^IFNγ^-/-^ mice (Fig. 3d,e). These data demonstrate that cytokines elicited by the immune response to infection disrupt stromal-nerve communication that is required to maintain sympathetic innervation of the spleen.

### Sympathetic nerve retraction modulates immunoregulatory functions in the spleen

Given the immunomodulatory role of sympathetic neurons, including via inflammatory reflexes that act in part via the spleen, we postulated that sympathetic anti-inflammatory functions would be reduced following nerve retraction. Because immune cells express adrenergic receptors and can respond to noradrenaline released from sympathetic nerves^17^, we first tested if replenishing the reduced adrenergic signals could impact adaptive immune responses during infection. We infected mice with LCMV and treated daily for 7 days with the non-selective beta-adrenergic receptor agonist isoprenaline (Extended Data Fig. 4g). Consistent with our hypothesis, antiviral CD8^+^ and CD4^+^ T cell responses were significantly reduced 8 days after infection in treated mice. We next tested if the capacity of sympathetic neurons to regulate systemic inflammation was impaired following spleen nerve retraction. Sympathetic neurons regulate acute inflammation following systemic challenge with lipopolysaccharide (LPS) by limiting the production of cytokines including TNF^6,34^. We used chemogenetic mice expressing an excitatory DREADD (hM3Dq) in TH-expressing neurons (TH-hM3Dq) and activated sympathetic neurons in these mice using the ligand clozapine N-oxide (CNO)^16^. Naïve and LCMV infected mice were treated with CNO and then administered LPS before TNF production was measured in serum 1.5 h later (Fig. 3i). As anticipated^1^, chemogenetic activation of TH-hM3Dq mice inhibited TNF production in uninfected mice following LPS challenge (Fig. 3j). In contrast, chemogenetic activation was unable to significantly decrease inflammation in infected mice.

These data suggested that the capacity of sympathetic neurons to modulate inflammation was blunted after infection, correlating with retraction of spleen innervation. To investigate splenic responses, we next performed intrasplenic injections of a recombinant AAV vector carrying a Cre-dependent hM3Dq-mCherry in TH-Cre mice before infection with LCMV 3 weeks later (Fig. 3k). This neuronal targeting approach transduced neuron somata in the SCSMG in Cre^+^ mice, which were visualised by expression of mCherry (Fig. 3l). DREADD expression was not observed after intrasplenic AAV injection into Cre^-^ mice. Chemogenetic activation of targeted neurons in naïve mice resulted in an inhibition of pro-inflammatory IFNγ and TNF cytokine responses in the spleen, while increasing anti-inflammatory IL-10 production (Fig. 3m and Extended Data Fig. 4h). In contrast, the levels of splenic pro-inflammatory and anti-inflammatory cytokines were not significantly modulated by chemogenetic activation in LCMV infected animals. In response to LPS challenge, NK cells and NKT cells are the main early producers of IFNγ in the spleen^35^. Chemogenetic activation reduced the production of IFNγ by these cells in the spleens of uninfected mice but not infected mice (Extended Data Fig. 4i). Together, these results reveal that following spleen nerve retraction that is induced by infection, the sympathetic inflammatory control in the spleen is functionally diminished.

### Infection induces inflammation of sympathetic ganglia

Loss of sympathetic nerve axons in the spleen after infection led us to question whether the neuronal somas located in the prevertebral ganglia were impacted. Histological examination of ganglia did not reveal any virus in the SCSMG, nor any obvious reduction in neuronal cell density (Extended Data Fig. 5a, b). Spleen innervation was restored 28 days after infection (Fig. 2e), further signifying that the neuron cell bodies did not die. To investigate impacts on SCSMG neurons during infection, we examined snRNA-seq of whole SCSMG isolated from LCMV infected mice (Fig. 4a). This revealed a proportional reduction in glial cells and the appearance of lymphocytes and myeloid cells in the ganglia tissue after infection (Fig. 4b). Differential gene expression analysis showed that neurons and glial cells responded robustly during infection by regulating hundreds of genes (Fig. 4c). This included upregulation of genes involved in anti-viral immune responses, antigen presentation pathway genes and interferon responsive genes (Extended Data Fig. 5c,d). Sympathetic neurons increased expression of MHC molecules and antigen processing pathway genes including *B2m, H2-K1* and *Psmb8*, as well as interferon-stimulated genes, including *Oasl2*, *Ifi27l2a* and *Irgm2* (Fig. 4d, e). We performed flow cytometry on dissociated SCSMG from TH-tdTomato mice, which confirmed that Tomato^+^ neurons upregulated surface MHC-I expression after infection, demonstrating neuronal responses during infection (Fig. 4f and Extended Data Fig. 5e). Infection also induced an increase in the chemokine receptor *Cxcr4*, the RNA binding protein HuD (*Elavl4*) and the neuroprotective gene *Oxct1* (Fig. 4e). The regulatory beta subunit of BK ion channels, *Kcnmb4*, which has been implicated in control of neuron excitability^36^, was also upregulated after infection. An increase in antigen presentation and interferon-stimulated genes was also observed in SCSMG glial cells (Extended Data Fig. 5f). Thus, LCMV infection caused inflammation of the sympathetic prevertebral ganglia that involved inflammatory responses by neurons and glial cells.

**Figure 4:**
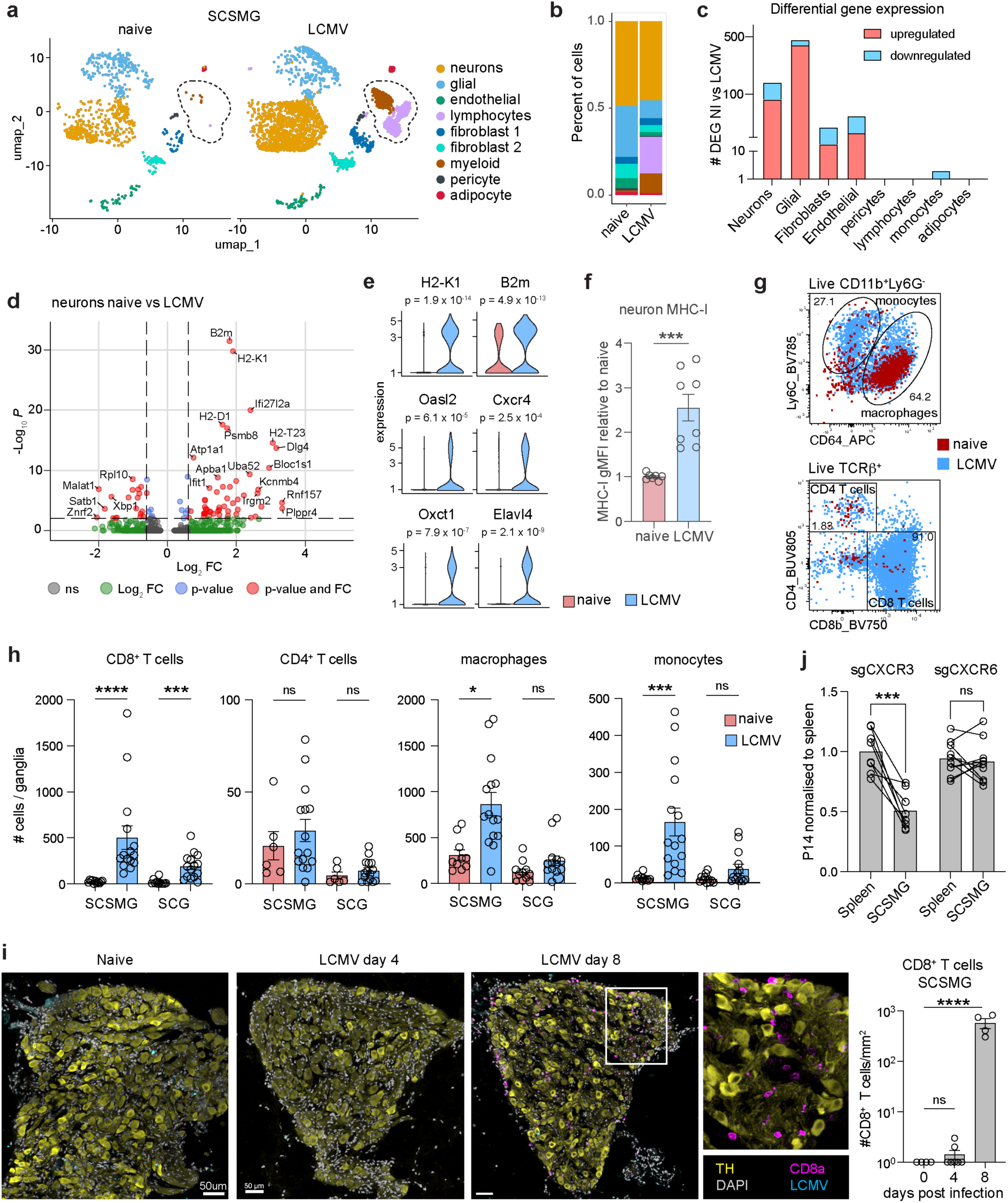
Systemic infection induces neuroinflammation and immune cell recruitment to sympathetic ganglia. **a**) snRNA-seq analysis of mouse SCSMG from naïve (757 cells) and LCMV (2983 cells) infected mice. Immune cell clusters are highlighted by the dotted regions. **b**) Cluster proportions in the naïve and LCMV SCSMG samples. **c**) Differential gene expression analysis comparing numbers of up or downregulated genes in SCSMG neurons from naïve and LCMV infected mice. **d**) Volcano plot of DE genes in neurons from naïve and LCMV mice. **e**) Violin plots of selected genes upregulated in SCSMG neurons after LCMV infection. *P* values from differential gene expression analysis are shown above the plots. **f**) Flow cytometry analysis of TH-tdTomato^+^ neurons from SCSMG from naïve and day 8 LCMV mice stained for surface MHC-I expression. **g, h**) Assessment of immune cell infiltration into SCSMG and SCG after LCMV infection. **g**, Example flow cytometry plots with naïve and LCMV samples overlaid. **h**, enumeration of CD8^+^ and CD4^+^ T cells, macrophages and monocytes. n = 11-15 mice per group. **i**) Imaging of SCSMG from naïve mice and mice 4 or 8 days after LCMV infection. Sections were stained for TH, CD8a, DAPI and for LCMV (not detected). Numbers of CD8^+^ T cells were quantitated from tissue sections (graph). n = 4 mice per group. **j**) Control (sg*CD19*) and CXCR3 (sg*CXCR3*) or CXCR6 (*sgCXCR6*) deficient P14 T cells were co-transferred into recipient mice prior to LCMV infection; spleen and SCSMG analysed 8 d post-infection. Ratios of P14 cells are normalised to spleen. Graphs show data (mean ± SEM) from 2-3 independent experiments. *P < 0.05, ***P < 0.001, ****P < 0.0001, ns, non-significant; by Mann-Whitney test (f), Kruskal-Wallis test (h), paired t-test (j) or one-way ANOVA (i).

### Recruitment of immune cells into inflamed ganglia

Given the inflammatory gene expression observed in SCSMG after infection, we examined the immune cell clusters found by snRNA-seq. Subsets of lymphocytes, including central memory and effector phenotype CD4^+^ and CD8^+^ T cells, as well as macrophages and monocytes, were present in the ganglia after infection (Extended Data Fig. 5g, h). Investigation of immune cell recruitment to the SCSMG was conducted by performing flow cytometry on dissociated ganglia tissues (Fig. 4g and Extended Data Fig. 6a). This revealed a significant recruitment of CD8^+^ T cells, but not CD4^+^ T cells, to the SCSMG after infection (Fig. 4h). CD8^+^ T cells were also recruited to the SCG after infection, albeit in lower numbers when compared to SCSMG. Macrophages and recruited monocytes were significantly increased in number in the SCSMG after infection but were not significantly increased in the SCG, indicating responses were focused on the ganglia that connect to the spleen (Fig. 4h).

Examination of SCSMG by confocal microscopy showed that recruitment of T cells occurred after day 4 of infection (Fig. 4i), which was after the initial reduction in innervation in the spleen. CD8^+^ T cells were observed distributed throughout the ganglia tissue, often in close proximity to TH^+^ neurons (Fig. 4i, magnified region). Transfer of LCMV-specific P14 CD8^+^ T cells prior to infection revealed that virus-specific effector T cells were recruited to the ganglia after infection (Extended Data Fig. 6b). These cells were phenotypically distinguishable from effector T cells in the spleen; a proportion of the cells upregulated CD69 and downregulated CX3CR1 in the ganglia (Extended Data Fig. 6c). Since this phenotypic change potentially reflected the development of a tissue resident phenotype by the recruited T cells, we examined mice 30 days after LCMV infection. This revealed that some anti-viral T cells persisted in the SCSMG during the memory phase and predominantly expressed a resident CD69^+^CX3CR1^-^ phenotype. Finally, we investigated the mechanism of CD8^+^ T cell recruitment to SCSMG. T cells can use CXCR3 and CXCR6 to enter and accumulate in the central nervous system^37,38^. We found increased expression of the CXCR3 ligands *Cxcl9* and *Cxcl10* by infiltrated myeloid cells, endothelial cells and pericytes after infection, as well as increased expression of *Cxcl16*, the ligand for CXCR6, by myeloid cells and pericytes (Extended Data Fig. 6d). We knocked down expression of *Cxcr3* or *Cxcr6* in P14 CD8^+^ T cells (Extended Data Fig. 6e), which demonstrated a requirement for CXCR3 but not CXCR6 for the recruitment of T cells into SCSMG after infection (Fig. 4j). Together, these results show that infection precipitated neuroinflammation in SCSMG, marked by inflammatory responses by sympathetic neurons and glial cells, and the subsequent recruitment of immune cells.

### Immune responses temper sympathetic neuron functions

Rather than showing spontaneous activity or behaving as ‘integrate and fire’ neurons, sympathetic ganglion cells typically relay strongly suprathreshold preganglionic inputs in a 1:1 fashion^39^. However, they also receive ongoing weaker synaptic inputs, whose chance of reaching threshold and leading to postganglionic firing can be significantly reduced by modest changes in neuron excitability^40^. Therefore, we sought to directly investigate whether virus infection alters the function of sympathetic neurons. To do this, we isolated SCSMG from naïve and LCMV infected mice and compared the characteristics of single neurons within intact ganglia using intracellular microelectrode electrophysiology^41^ (Fig. 5a). We determined that the minimum electrical current required to elicit action potentials (the rheobase) in SCSMG sympathetic neurons was significantly higher after infection when compared to uninfected mice (Fig. 5b,c). The resting membrane potential of sympathetic neurons was unchanged by infection; however, the cells’ input resistance was significantly reduced, indicating that leak currents were increased during infection (Fig. 5c). We detected these changes in all neurons randomly selected in SCSMG, signifying that the functional changes were not limited to spleen-innervating neurons. We questioned if the altered neuronal excitability in the SCSMG resulted from the inflammatory recruitment of T cells into the ganglia. However, assessment of neuronal excitability 3-4 days after LCMV infection, when spleen innervation was retracting (Fig. 2e), and prior to T cell infiltration into SCSMG (Fig. 4i), revealed that neuronal hypo-responsiveness was already induced (3.8-fold higher than uninfected mice) (Fig. 5c). Furthermore, although T cells were recruited to the SCG, the excitability of neurons in the SCG was not changed after infection (Fig. 5d and Extended Data Fig. 7a). Thus, sympathetic hypo-excitability was limited to neurons in the ganglia that connect to the spleen, indicating that circulating inflammatory stimuli were insufficient to downregulate neuronal function in distant ganglia. Within 4 weeks of infection neuron firing threshold and input resistance had returned to baseline (Fig. 5c). This correlated with re-innervation of the spleen (Fig. 2e) and signified that neuronal hypo-excitability induced after infection was reversible.

**Figure 5:**
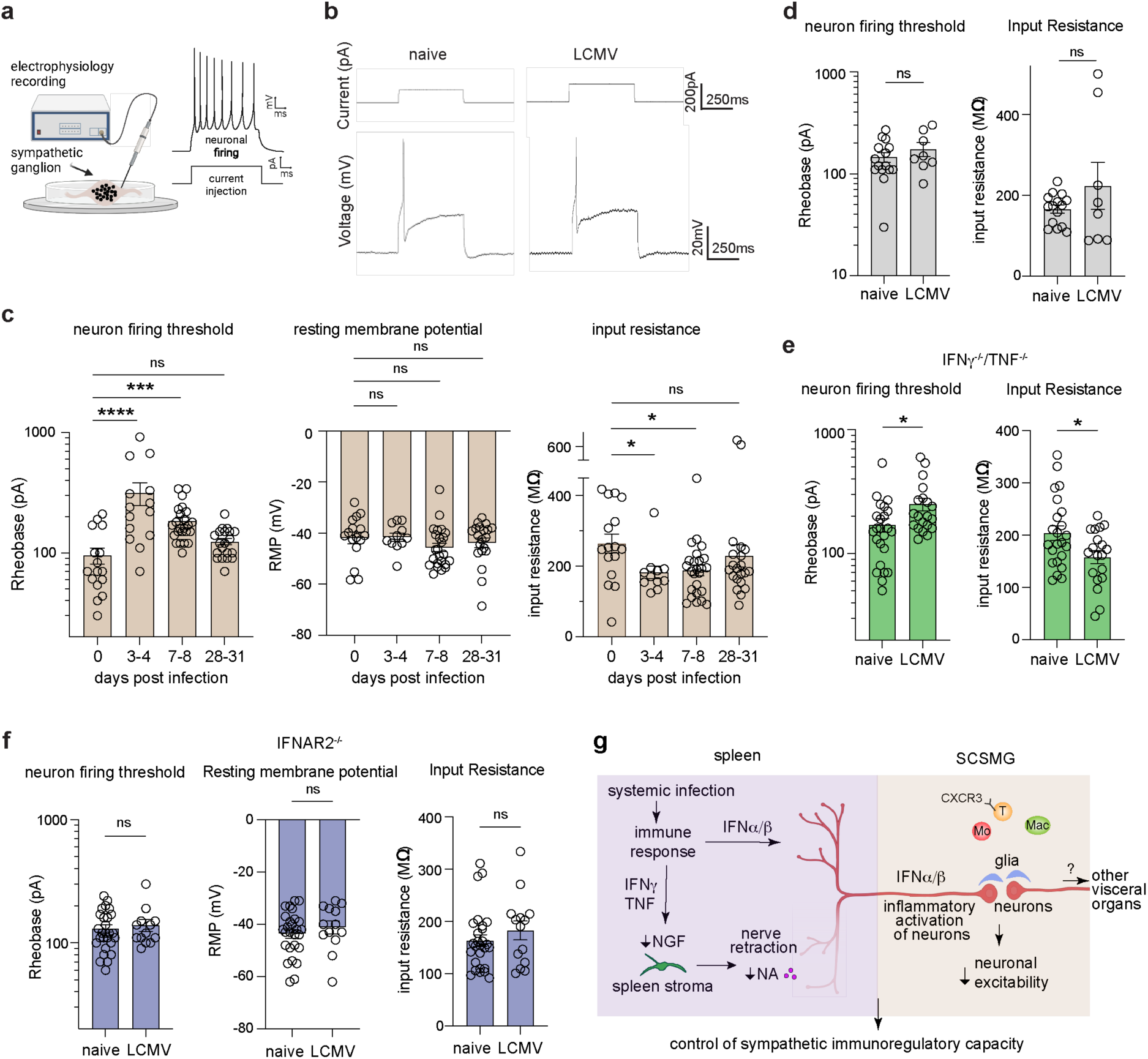
Splenic type I interferon signalling selectively impairs sympathetic neuron excitability in the SCSMG during infection. **a**) Microelectrode intracellular electrophysiology recordings from intact SCSMG was performed in naïve mice and after LCMV infection to determine sympathetic neuron electrical characteristics. **b**) Representative depolarizing current step (top trace) required to elicit an action potential recorded from a neuron in the SCSMG (bottom trace). Microelectrodes inserted into single SCSMG neurons were used to record the electrical activity. **c**) Neuron firing thresholds, resting membrane potentials and input resistance in SCSMG from naïve (day 0) and LCMV infected mice (days 3-4, 7-8 and 28-31). n = 11-24 neurons from 4-8 mice. **d**) Neuron firing thresholds and input resistance recorded in SCG from naïve and day 8 LCMV mice. n = 8-15 neurons from 3 mice. **e**) Neuron firing thresholds and input resistance recorded in SCSMG from IFNγ^-/-^/TNF^-/-^ mice. n = 20-24 neurons from 4 mice. **f**) Neuron firing thresholds, resting membrane potentials and input resistance in SCSMG from IFNAR2^-/-^ mice. n = 13-21 neurons from 4 mice. Graphs show representative data (mean ± SEM) from 2-3 independent experiments. *P < 0.05, **P < 0.01, and ***P < 0.001, ns, non-significant; by one-way ANOVA (c) or unpaired t-test (d-f). **g**) Proposed model of the regulation of sympathetic neuron functions during infection. Infection of the spleen involves the production of TNF and IFNγ by immune cells that include NK cells and T cells and leads to a reduction in NGF expression, loss of spleen TH^+^ sympathetic innervation and reduced spleen noradrenaline. Concurrently, IFNα/β signals selectively impair neuronal excitability in the SCSMG, but not SCG. The reduced sympathetic neuron functions modulate sympathetic immunoregulatory capacity in the spleen, which could support immunity. Effects of neuron hypo-excitability on other organs is currently unknown.

To further investigate the mechanism of reduced neuron excitability, we reasoned that TNF and IFNγ might be involved because these cytokines were responsible for loss of innervation in the spleen. However, SCSMG neurons from IFNγ^-/-^/TNF^-/-^ mice displayed significantly increased neuronal firing thresholds and reduced input resistance (Fig. 5e and Extended Data Fig. 7b), demonstrating hypo-excitability and suggesting that additional mechanisms were responsible for the changes in neuron function after infection. The early changes in neuron function, combined with interferon response signatures in SCSMG neurons (Fig. 4d, e) led us to examine a role for type I interferons. Electrophysiological recordings of sympathetic neurons in IFNAR2^-/-^ mice, which lack the receptor subunit essential for responding to type I interferons, revealed no decrease of neuronal excitability after LCMV infection (Fig. 5f). These data reveal a type I interferon-dependent functional impairment in sympathetic neurons in the SCSMG during pathogen infection. Together, our study reveals that the immune system modulates the sympathetic nervous system during the response to infection, via contemporaneous organ-specific retraction of spleen innervation and reduced neuronal function.

## Discussion

Peripheral sympathetic neurons act as an efferent arm of an endogenous inflammatory reflex, conveying signals from the brain that regulate inflammatory responses in the body^1,2,7^. The spleen is an important site where inflammatory responses are regulated by this sympathetic ‘brake’. Although sympathetic nerves in the spleen that produce noradrenaline and NPY have been shown to act on immune cells to suppress inflammation ^3,8,15^, the nature of these neurons and whether the immune system can reciprocally regulate peripheral sympathetic neurons has remained unknown. Here we identify a control mechanism that regulates the efferent inflammatory reflex via signals elicited during immune responses to infection. This neuroimmune pathway transiently controls splenic sympathetic innervation and neuronal functions to regulate sympathetic activity (Fig. 5g). We propose that this enables organ-specific control of sympathetic immunoregulation that is localised to the site of immune responses induction, supporting immunity.

Although it had been proposed from functional studies that a distinct class of sympathetic neuron that is specialised for communication with immune cells innervates lymphoid organs^20^, our transcriptional profiling of spleen-innervating neurons did not identify a distinct subset in the spleen. Nonetheless, we found that spleen sympathetic neurons are heterogeneous, suggesting spleen innervation may be more diverse than other visceral organs where recent work discovered that organ-specific innervation is restricted to only one of two molecular subtypes^27^. But few studies have characterised the transcriptome of sympathetic neurons where the innervated target organ is known^26,42^, leaving open the possibility that intra-organ sympathetic neuron heterogeneity is more diverse than we currently understand. Additional investigation of neuronal diversity will be required, including analysis of epigenetic mechanisms. In the spleen we found that the white pulp is innervated by RXFP1^+^ neurons, while the red pulp contains a heterogeneous population. It remains to be established whether this represents innervation of discrete spleen tissue niches or cell types by these transcriptionally heterogeneous neuron populations. But our data suggest that instead of a specific immunoregulatory sympathetic neuron class in the spleen, feedback between immune cells and post-ganglionic neurons facilitates fine tuning of neuroimmune circuits.

The spleen plays a central role in protecting the body against systemic infections through an organised microanatomy^43^. By revealing that the cytokines TNF and IFNγ act via a stromal cell-NGF axis to control spleen innervation, our data provides a new basis for understanding the complex interplay between the sympathetic nervous system and the immune system. Further studies are required to ascertain if sympathetic nerve retraction also shapes immunoregulation in lymph nodes or primary lymphoid tissues. Pathological changes in spleen sympathetic innervation have been observed during chronic infection with murine HIV^44^ and in patients that died from sepsis^45^. In contrast, we find that acute infection induces a rapid and transient nerve retraction, although it is possible that pathological conditions perturb the same stromal-NGF axis in the spleen. Unlike the sympathetic neuropathy observed in the SCG during cardiac disease^25^, we did not find death of neurons in the ganglia, which likely enabled innervation of the spleen to be restored within weeks. Our experiments also revealed retention of sensory nerves in the spleen during infection. Since sensory nerves can support T and B cell responses^11,12^, loss of sympathetic signals in the spleen could complement nociceptor-driven support for immune responses.

Our results also showed that sympathetic neurons were contemporaneously regulated by interferon signals to a hypo-responsive state. This change in neuron excitability was observed in SCSMG, ganglia with connections to the spleen, suggesting that interferon signals are received by axons in the spleen and retrogradely signal to the soma, but not in SCG neurons. Yet, all SCSMG responded to inflammatory signals and became hypo-responsive after infection, reflecting changes not only to spleen-innervating neurons but many other neurons that innervate different visceral organs. It will be interesting to determine if this hypo-responsiveness also modifies physiological sympathetic nervous system functions in other visceral organs, potentially impacting vasculature and gastrointestinal functions. Exploration of the broader impacts of immune responses on sympathetic nervous system functions has the potential to reveal new therapeutic targets for post-infectious sympathetic dysautonomias.

## Acknowledgements

This work was supported by grants from the National Health and Medical Research Council of Australia, grant 2017220 to S.N.M. and grant 1186384 to R.M.M.; P.T-B. receives the University of Florida McKnight Brain Institute Gator NeuroScholar Fellowship. We thank Martin Stebbing, Michael McKinley, Bill Heath, and Stuart Mazzone for discussions and advice. This research was supported by The University of Melbourne’s Research Computing Services and the Petascale Campus Initiative. We acknowledge the Biological Optical Microscopy Platform at the University of Melbourne for use of the confocal microscopy; the Melbourne Cytometry Platform (Doherty Institute node, University of Melbourne) for the use of their Flow Cytometry instruments and their technical assistance; and the MHTP Medical Genomics Facility and the South Australian Genomics Centre for sequencing.

## Author contributions

Conceptualization, P.T-B., M.T.R., R.M.M., J.B.F., S.N.M.

Methodology, P.T-B., M.T.R., M.D., S.C.M., J.C.M., R.M.M., S.N.M.

Investigation, P.T-B., M.T.R., M.D., S.C.M., J.K.L., S.D., A.W., Y.O.A., D.S., H.L.H., Z.F., T.N.B., E.A.C., L.K., I.F., M.R.D.N., M.d.L.M,

Analysis, P.T-B., M.T.R., M.D., S.C.M., J.K.L., S.D., Z.L., A.W., L.D., H.J.L., S.L., D.R., S.N.M.

Resources, G.T.D., K.K., L.K.M., A.H.

Writing – Original Draft, P.T-B., S.N.M.

Writing – Review & Editing, all authors

Visualization, M.T.R., S.N.M.

Supervision, E.K.S., J.S., R.M.M., J.B.F., S.N.M.

Funding Acquisition, R.M.M., J.B.F., S.N.M.

## Declaration of interests

The authors declare no competing interests.

## Extended Data Figures

**Extended data Figure 1:**
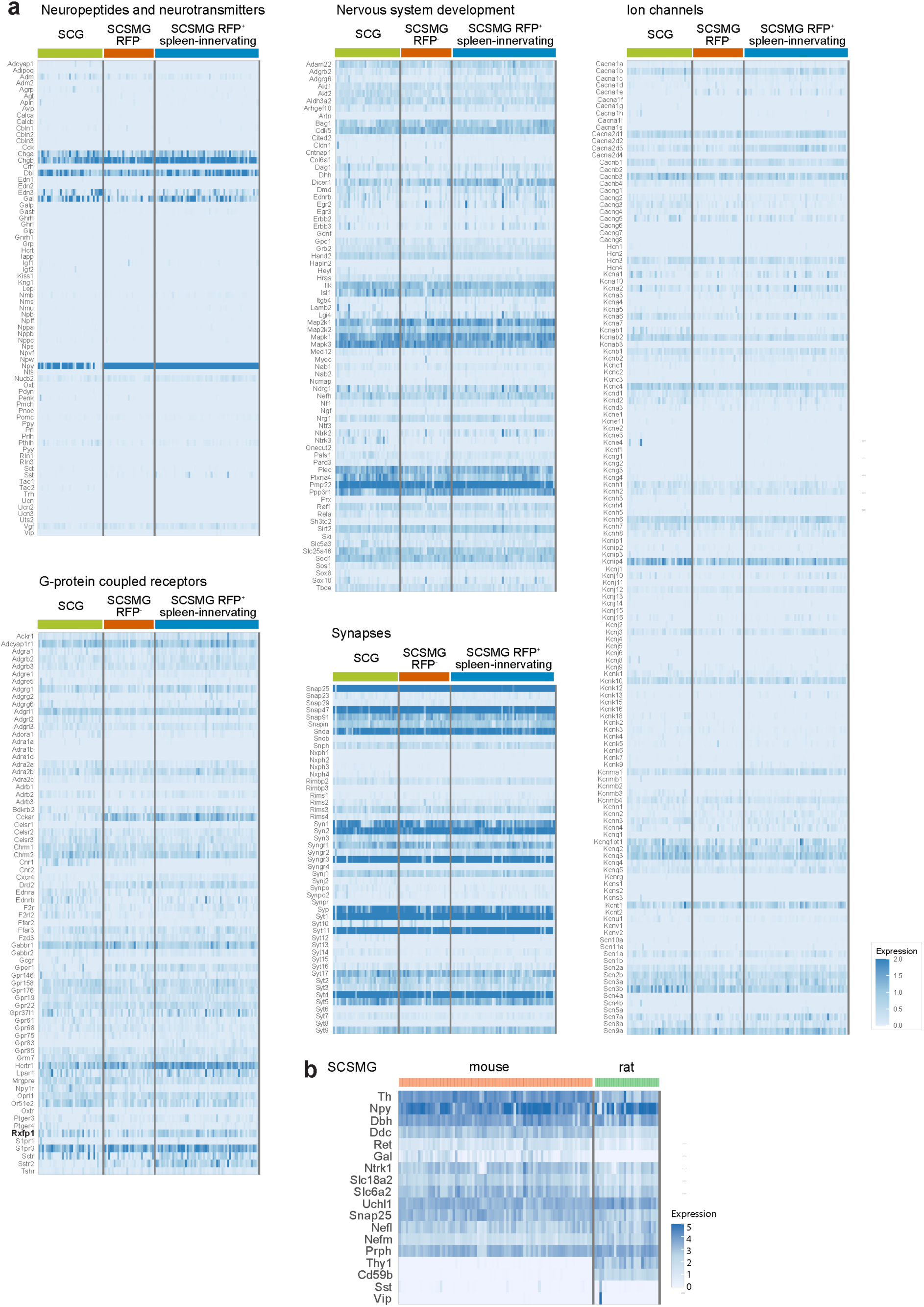
Transcriptomic analysis of mouse and rat spleen-innervating sympathetic neurons. **a**) Heatmaps depicting expression of neurotransmitter, synapse, nervous system development and ion channel genes amongst SCG, unlabelled (RFP^-^) SCSMG and labelled (RFP^+^) SCSMG neurons profiled by scRNA-seq. **b**) Heatmap of selected neuronal marker genes amongst 79 mouse SCSMG neurons and 26 rat SCSMG neurons profiled by scRNA-seq.

**Extended data Figure 2:**
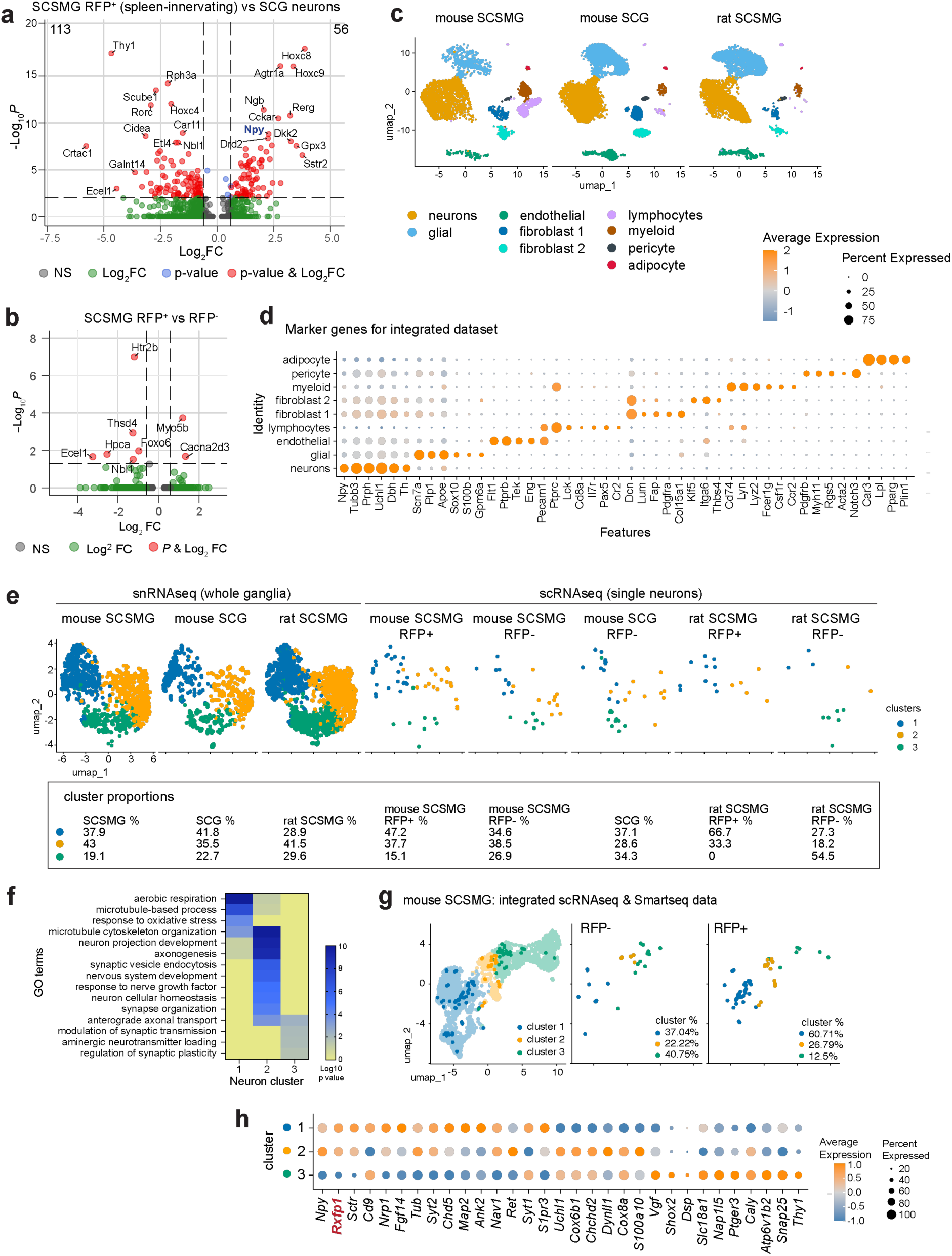
Multi-modal transcriptomic analysis of sympathetic neuron diversity. **a**) Volcano plot of differentially expressed (DE) genes in spleen-innervating neurons compared to SCG neurons. Selected genes are labelled, and numbers in upper corners represent the total number of DE genes. **b**) Volcano plot of DE genes in spleen-innervating RFP^+^ neurons compared to unlabelled SCSMG neurons. **c**) Split UMAP showing cluster distribution across the three sample groups. Mouse SCSMG: 3740 cells, mouse SCG: 7778 cells, rat SCSMG 4792 cells. **d**) Marker gene expression by cell type clusters from mouse SCSMG and SCG, and rat SCSMG analysed by snRNA-seq. **e**) UMAP projections of integrated snRNA-seq and scRNA-seq analysis of mouse and rat ganglia. The proportion of cells in each cluster is shown below the plots. **f**) GO term enrichment in the indicated neuron clusters. **g, h**) Integration of mouse SCSMG neurons profiled by Smartseq (119 cells) and profiled 10X scRNAseq (2466 cells) from ^26^. **g**) UMAP projections of integrated scRNA-seq analysis. The proportion of cells in each cluster is shown on the plots. **h**) DE genes amongst the three neuron clusters.

**Extended data Figure 3:**
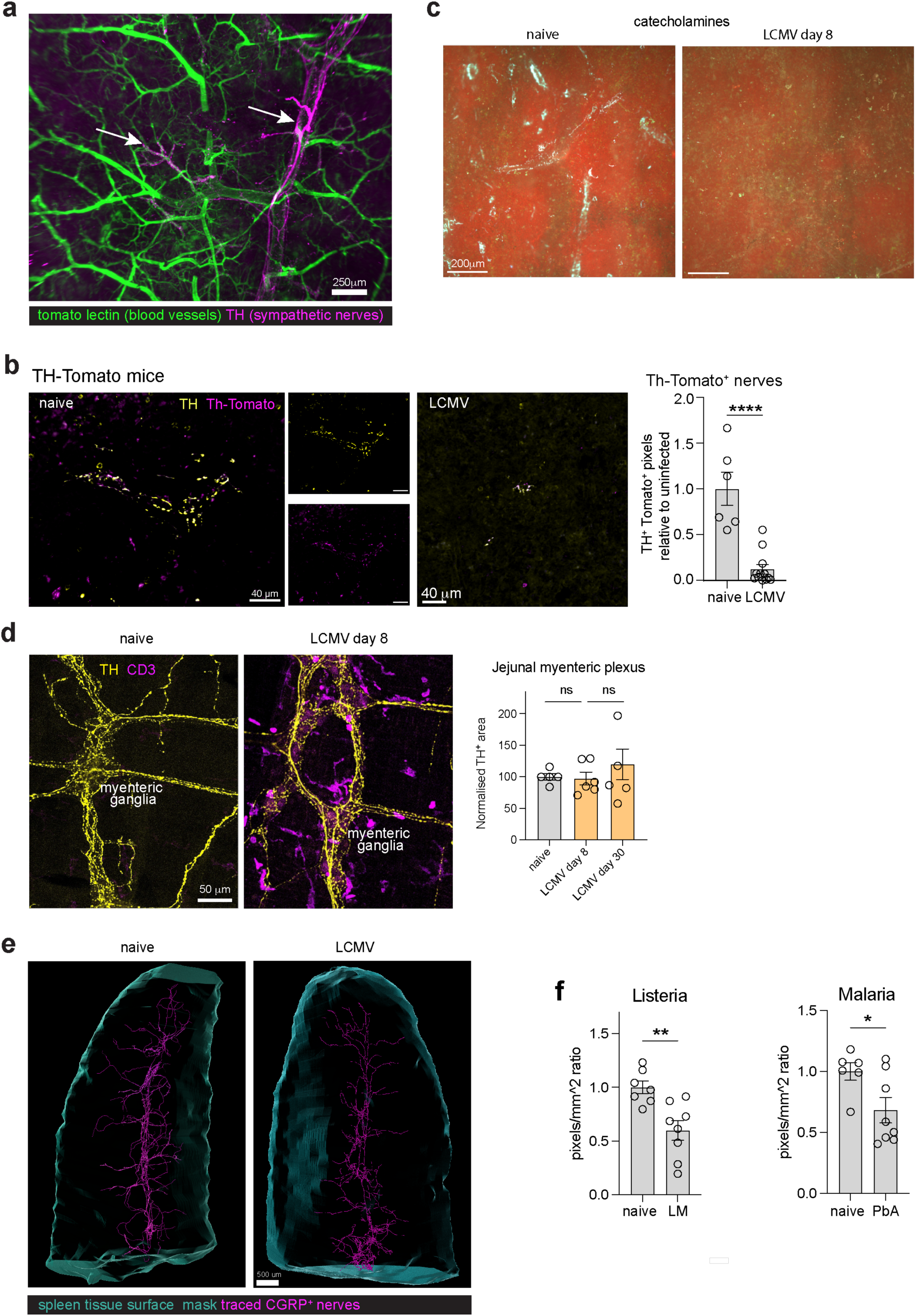
Retraction of spleen innervation after infection. **a**) Optical section (500 μm) of whole-mount spleen from day 8 LCMV mice that received an intracardiac injection of tomato lectin to label blood vessels and stained for TH prior to clearing and light sheet imaging. Arrows depict TH^+^ nerves that are retained on the main artery in the splenic hilum and occasional adjacent branches. **b**) Spleen innervation in TH-tdTomato mice stained with anti-TH antibody (yellow) (left) and quantitation of innervation in naïve and LCMV infected mice (right). n = 6-12, 2-3 regions each from 5-6 mice per group. **c**) Images of SPG staining of spleen sections from naïve mice and 8 days after LCMV infection to detect catecholamines (cyan staining). **d**) Whole-mount imaging of the small intestinal myenteric plexus from naïve and day 8 LCMV mice stained for TH and CD3. Location of myenteric ganglia (determined by HuC/D staining, not shown) is indicated. Quantitation of TH^+^ nerve fibres is graphed (right). n = 5-6 mice from 4 independent experiments. **e**) 3D views of CGRP^+^ sensory innervation (magenta) in the spleen in naïve and day 8 LCMV mice that was traced using Imaris. Spleen surface rendering is shown in cyan. **f**) Quantitation of sympathetic innervation in spleen sections from naïve and *Listeria monocytogenes* or *Plasmodium berghei* ANKA infected mice. n = 6-8 mice per group. Graphs show representative data (mean ± SEM) from 2-4 independent experiments. ****P < 0.0001, ns, non-significant; unpaired t-test (b, f) or by one-way ANOVA (d).

**Extended data Figure 4:**
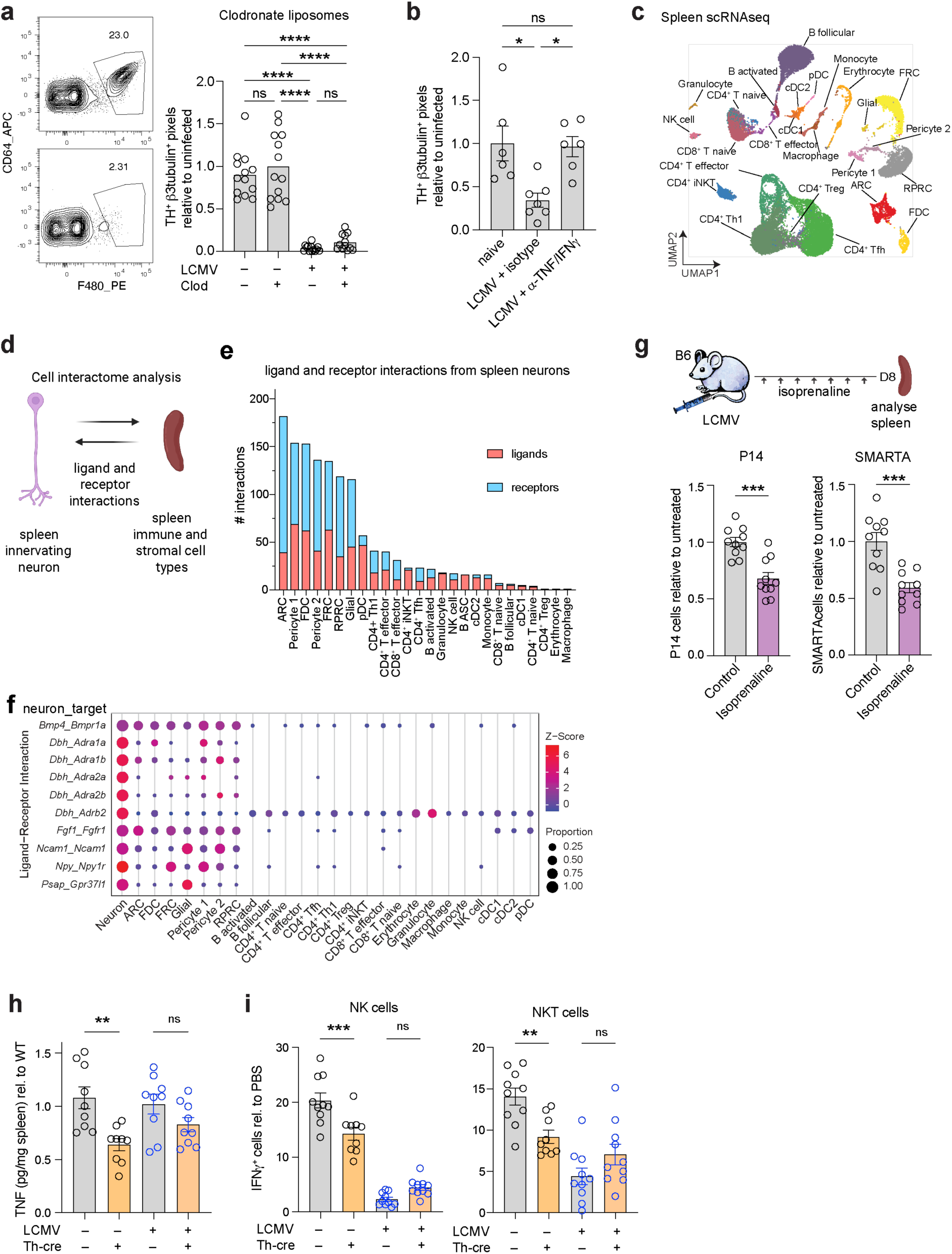
Splenic sympathetic nerve retraction modulates sympathetic control of inflammation. **a**) Clodronate liposome treatment of mice to deplete macrophages and infection with LCMV. Enumeration of spleen innervation in naïve and LCMV infected mice with or without clodronate liposome treatment. n = 8-9 mice per group. **b**) Quantitation of sympathetic innervation in spleen sections from naïve and LCMV infected mice treated with anti-TNF and anti-IFNγ blocking antibodies. n = 6-7 mice per group. **c**) UMAP of scRNA-seq data of spleen stromal and immune cells, from references^28,29^. **d**) Schematic depicting the interactome analysis that was performed to predict receptor-ligand interactions between mouse spleen-innervating neurons (RFP^+^) and subsets of mouse spleen stromal and immune cells. **e**) Number of potential ligand and receptor interactions from neurons to spleen cells, stratified by spleen cell type with the highest to lowest number of predicted total interactions. **f**) Top interactions predicted between spleen neurons and spleen stromal and immune cells. Shown are ligand-receptor interactions from spleen neurons to predicted interacting molecules on the indicated spleen cell subsets. ARC, adventitial reticular cell; FDC, follicular dendritic cell; FRC, fibroblastic reticular cell; RPRC, red pulp reticular cell. **g**) Reduction in transgenic P14 and SMARTA T cell responses to LCMV infection in the spleen in response to isoprenaline administration. Mice were administered isoprenaline daily from day 0-7 of infection and analysed on day 8. n = 10 mice per group. **h**) Intrasplenic injections of rAAV-hM3Dq-mCherry were performed in TH-Cre^+^ or Cre^-^ mice 17d before LCMV infection, followed by CNO administration and LPS treatment. TNF was measured in spleen supernatant 90 min after LPS administration. **i**) Intracellular IFNγ staining of NK cells and NKT cells from the spleens of mice from (h). n = 9-10 mice per group. Graphs show representative data (mean ± SEM) from two independent experiments (a-b, g, h-i). *P < 0.05, **P < 0.01, ***P < 0.001, and ****P < 0.0001, ns, non-significant; by one-way ANOVA (a-b, h-i) or unpaired t-test (g).

**Extended data Figure 5:**
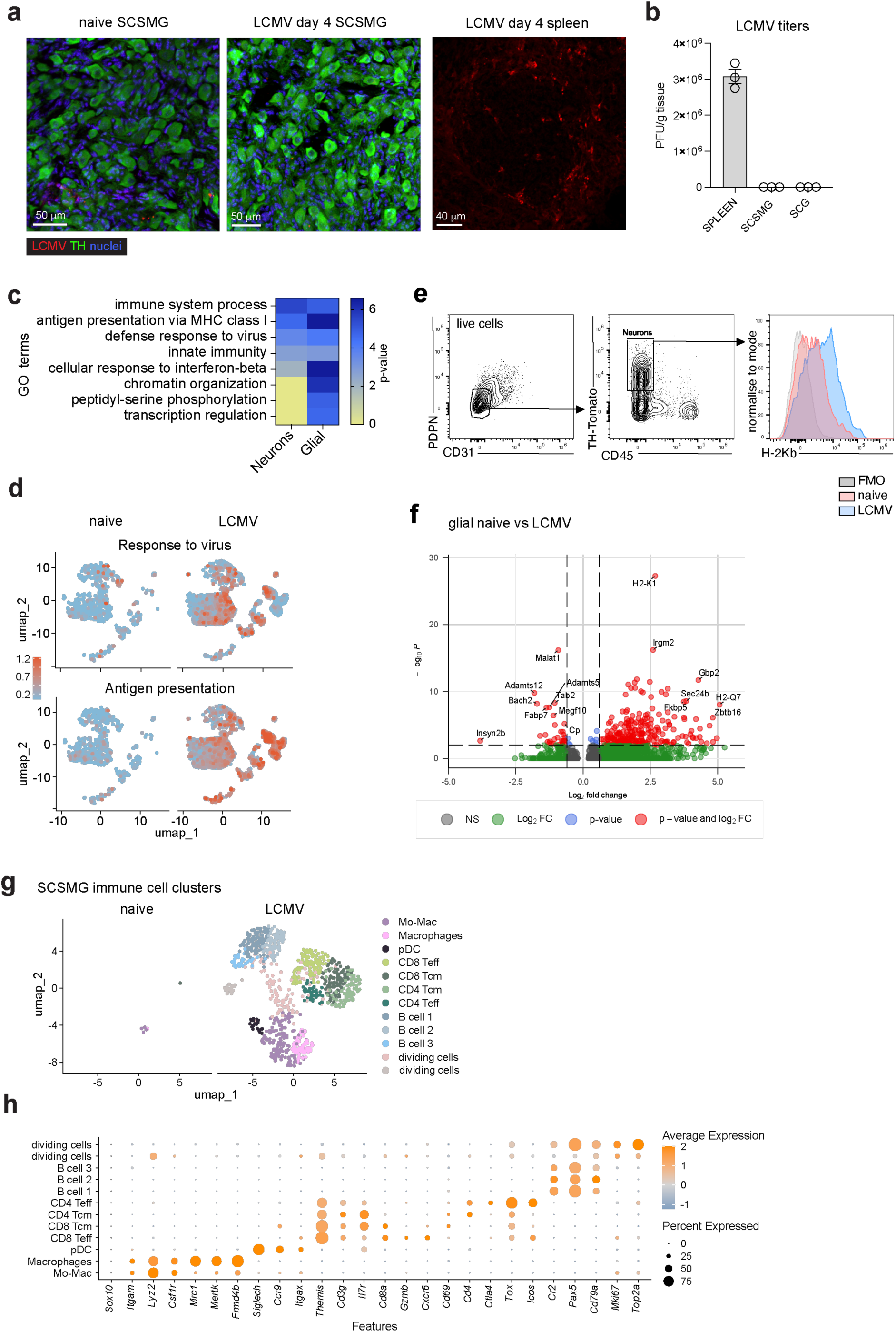
Neuroinflammation of SCSMG after infection. **a**) Imaging of SCSMG from naïve mice and day 4 LCMV mice, stained for TH (green), LCMV (red) and DAPI (blue), and day 4 spleen stained for LCMV. **b**) Viral titers in spleen, SCSMG and SCG 4 days after LCMV infection measured by pfu assay. n = 3 mice from one of two experiments. **c**) GO term analysis of DEG in neurons and glial cells between naïve and LCMV SCSMG. **d**) Expression of virus response and antigen presentation gene modules in naïve and LCMV SCSMG (see methods for details). **e**) Flow cytometry gating for TH-tdTomato^+^ neurons from the SCSMG of naïve and day 8 LCMV mice. **f**) Volcano plot of DE genes in glial cells from naïve and LCMV mice. **g**) UMAP comparing immune cells in SCSMG from naïve and LCMV infected mice by snRNA-seq. **h**) Marker gene expression by immune cell types from mouse SCSMG analysed by snRNA-seq.

**Extended data Figure 6:**
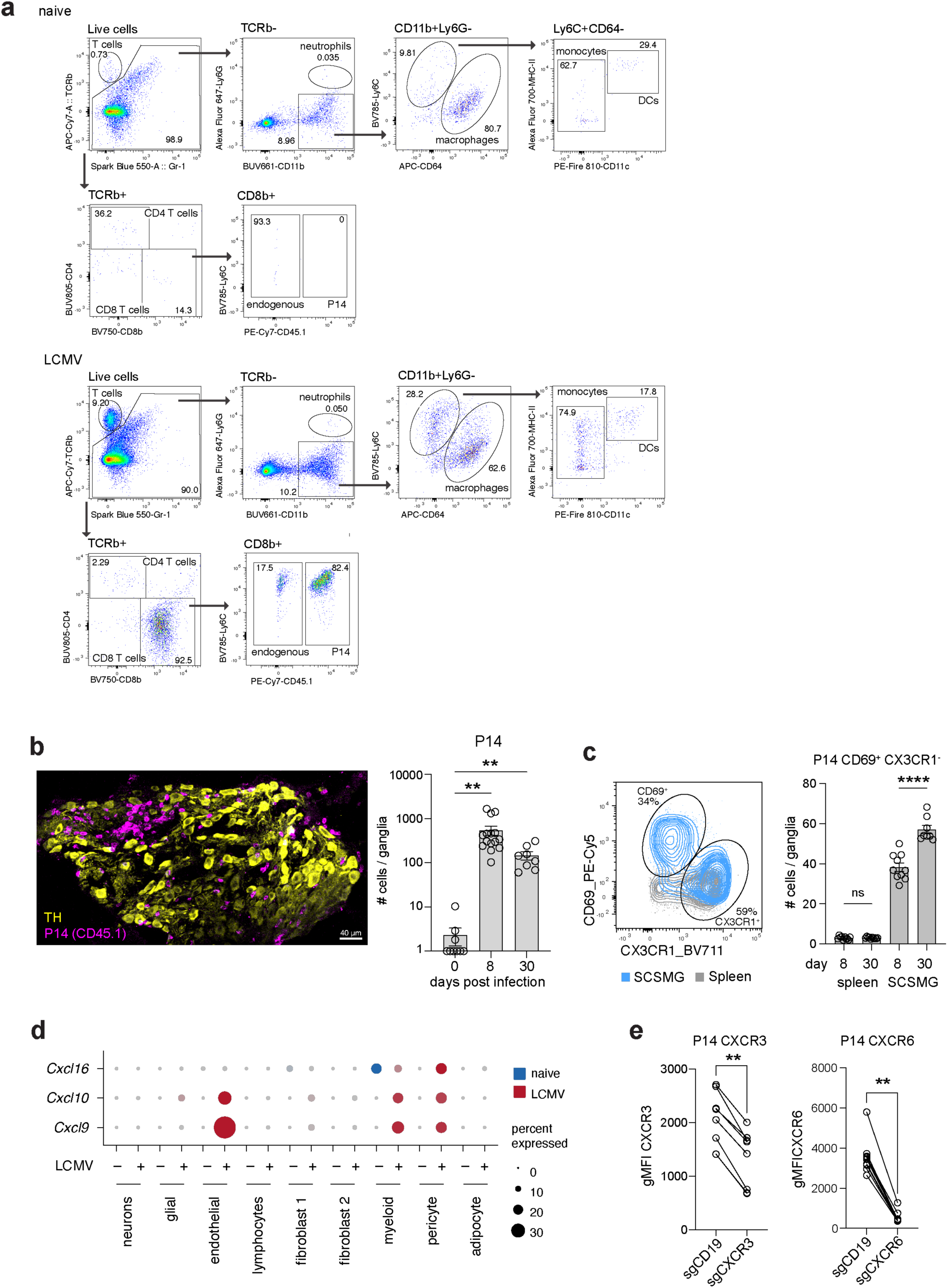
Recruitment and persistence of anti-viral T cells into inflamed sympathetic ganglia. **a**) Representative flow cytometry gating in naïve and day 8 LCMV infected mice to identify immune cell subtypes. **b**, **c**) Mice received P14.CD45.1 CD8 T cells prior to LCMV infection and SCSMG tissues were obtained 8-30 days later. **b**, Representative image of SCSMG from day 8 LCMV stained for TH and CD45.1 (left). Graph (right) shows enumeration of P14 CD8 T cells in SCSMG in naïve and LCMV infected mice 8 and 30 days after infection. n = 8-15 mice per group. **c**) The phenotype of P14 CD8 T cells isolated from spleen and SCSMG 8 days after infection (left) and the proportion of CD69^+^CX3CR1^-^ P14 T cells in the spleen and SCSMG 8 or 30 days after infection (right). **d**) Dotplot of snRNAseq data from Fig. 4a showing expression of *Cxcl16*, *Cxcl9* and *Cxcl10* on the indicated cell clusters in the SCSMG from naïve and day 8 LCMV mice. **e**) Expression of CXCR3 and CXCR6 on spleen P14 CD8^+^ T cells from day 8 LCMV mice. T cells were transduced with the indicated guide RNAs (sgCD19, control or sgCXCR3 or sgCXCR6) prior to adoptive transfer and infection. Graphs show representative data (mean ± SEM) from 2-3 independent experiments. **P < 0.01, and ****P < 0.0001, ns, non-significant; by Brown-Forsythe & Welch ANOVA (b-c) or paired t-test (e).

**Extended data Figure 7:**
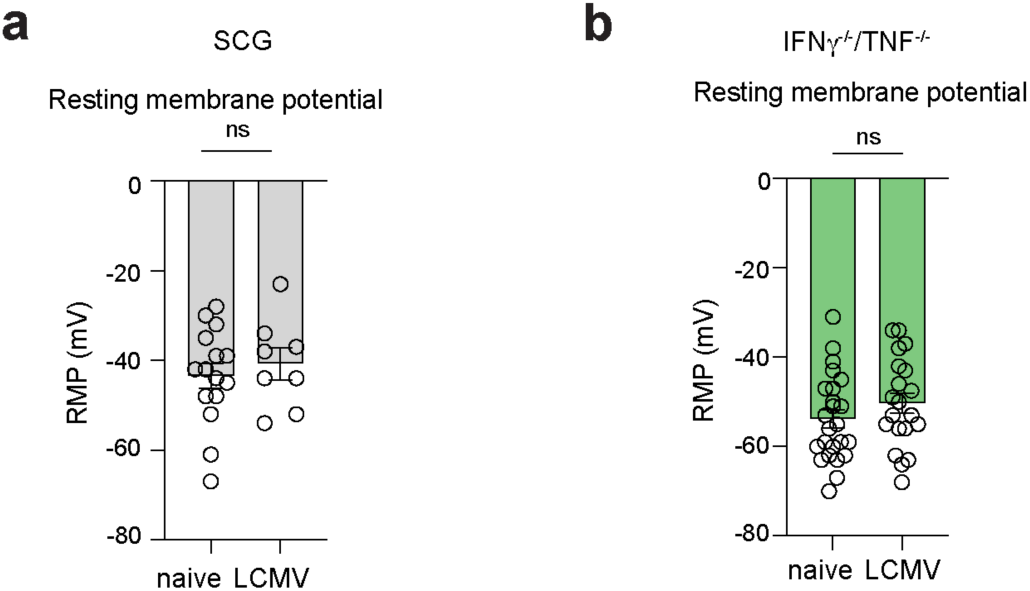
Modulation of sympathetic neuron functions during infection. **a**) Neuron resting membrane potentials and input resistance recorded in SCG from naïve and day 8 LCMV mice. n = 8-15 neurons from 3 mice. **b**) Neuron resting membrane potentials recorded in SCSMG from naïve and day 8 LCMV IFNγ^-/-^/TNF^-/-^ mice. n = 20-24 neurons from 4 mice. Graphs show representative data (mean ± SEM) from 2 independent experiments. ns, non-significant; by unpaired t-tests.

## Supplementary information

**Supplementary Video 1**

3D view of sympathetic nerves in healthy naïve spleen. Light sheet imaging of whole-mount spleen cleared using the Shanel method and immunolabelled with TH (magenta). Imaris surface rendering of the spleen surface is shown in cyan.

**Supplemental Video 2**

3D view of sympathetic nerves in an LCMV infected spleen. Light sheet imaging of whole-mount spleen cleared using the Shanel method and immunolabelled with TH (magenta). Imaris surface rendering of the spleen surface is shown in cyan.

**Supplemental Video 3**

3D view of sensory nerves in a naïve spleen. Light sheet imaging of whole-mount spleen cleared using the Shanel method and immunolabelled with CGRP to stain for sensory nerves. Imaris tracing of these CGRP sensory nerves is shown in magenta, whereas surface rendering of the spleen surface (using Imaris) is shown in cyan.

**Supplemental Video 4**

3D view of sensory nerves in a LCMV infected spleen. Light sheet imaging of whole-mount spleen cleared using the Shanel method and immunolabelled with CGRP to stain for sensory nerves. Imaris tracing of these CGRP sensory nerves is shown in magenta, whereas surface rendering of the spleen surface (using Imaris) is shown in cyan.

## Methods

### Animals and infections

C57BL/6 (B6), TNF^-/-^ (B6.129S-*Tnf^tm1Gkl^*/J), IFNγ^-/-^ (B6.129S7-*Ifng^tm1Ts^*/J), IFNAR2^-/-^ (C57BL/6-*Ifnar2^tm1Pjh^*), P14 x B6.SJL-*Ptprc^a^ Pepc^b^*/BoyJ (P14.CD45.1), P14/uGFP/CD45.1, B6.SMARTA/uGFP, CCL19-Cre^46^, B6.Cg-*Gt(ROSA)26Sor^tm14(CAG–tdTomato)Hze^*/J (Ai14), B6.Cg-*7630403G23Rik^Tg(Th–cre)1Tmd^*/J (Th-Cre)^47^, B6N;129-Tg(CAG-CHRM3*,-mCitrine)1Ute/J (R26-LSL-hM3Dq)^48^ and B6.Cg-Rxfp1^em1(cre)Ngai^/TasicJ (Rxfp1-Cre)^49^ mice were used. TH-Cre.R26-LSL-hM3Dq (TH-hM3Dq) designer receptors exclusively activated by designer drugs (DREADD) mice were validated previously^16^. Mice were bred at the Bioresource Facility of the Department of Microbiology and Immunology, The University of Melbourne. Mice were maintained under specific pathogen free conditions and housed in individually ventilated cages. Both female and male mice were used between 8-14 weeks of age. Mouse experiments were approved by the Animal Ethics Committee of the University of Melbourne. Mice were intraperitoneally infected with 2 x 10^5^ plaque-forming units (PFU) of LCMV Armstrong. Mice were adoptively transferred with 1-5 x 10^4^ P14 and SMARTA T cells prior to LCMV infection. To activate beta adrenergic receptors during LCMV infection, mice were treated daily from day 0-7 with i.p. injections of 5 mg/kg isoprenaline.

Sprague Dawley rats (body weight 250-400 g) were used. Rats were supplied by Animal Resources Centre, Perth, Western Australia. Rat experiments were approved by the Animal Experimentation Ethics Committee of the Florey Institute of Neuroscience and Mental Health and adhered to the guidelines of the National Health and Medical Research Council of Australia.

### Spleen retrobead microinjections

Initially, glass micropipettes for spleen microinjections were produced from Z114952-200EA capillary tubes (Sigma-Aldrich). The capillaries were pulled using a micropipette puller, generating a tip at around 80 µm. To make them sharp for the microinjections, the micropipettes were bevelled (45° angle). The micropipettes were connected to a 5 µl Hamilton syringe via silicone tubing, and mineral oil was backfilled into the system. The Hamilton syringe was placed on a single-syringe infusion pump (Aladdin WPI) and, just before performing the spleen microinjections, the micropipette was backfilled with Red Retrobeads^TM^ IX (Lumafluor, Inc.). For the surgery, mice were anaesthetised with i.p. ketamine (100 mg/kg body weight) and xylazine (15 mg/kg bodyweight); and rats were anaesthetized with i.p. ketamine (75 mg/kg body weight) and medetomidine (0.5 mg/kg body weight). The fur was clipped over the left side right under the rib cage. A skin incision (about 1 cm long in mice and about 2 cm long in rats) was made on the left side of the abdomen, just below the ribs, followed by a small incision in the muscle and peritoneum. Using sterile cotton swabs, the spleen was exposed externally and Red Retrobead microinjections were performed into the spleen. 100 nl of beads was injected per injection across several locations along the spleen. For the mice, a total of 2 µl of Red Retrobeads were injected; for the rats, 5 µl of Red Retrobeads were injected. The spleen was regularly moistened with PBS during the procedure to prevent drying. After performing microinjections in mice, we used sterile suture lines to close the muscle and peritoneum first, then the skin. In rats, the muscle and peritoneum were closed with sterile suture lines, and the skin was closed with fine wound stainless steel animal surgical clips. Animals were euthanized 7-12 days later to collect the ganglia for the localization of the retrogradely labelled neurons in the SCSMG.

### Ganglia isolation

Rats were sacrificed by i.p. injection of sodium pentobarbitone (100 mg/kg), and mice were humanely killed by cervical dislocation. SCSMG were isolated using our published protocol^23^. Briefly, a midline incision (in the Linea Alba) to open the abdominal wall was performed, and the abdominal viscera was pulled to the left with a sterile cotton swab. Under a dissecting microscope to magnify the surrounding region of the spleen and left kidney, the left splanchnic ganglion (SplG), the celiac ganglia (CG) and the superior mesenteric ganglion (SMG) were located. With fine forceps, the ganglia were dissected. Then, the SCSMG were transferred onto a Sylgard coated dissection dish with cold PBS. The SCSMG were pinned down onto the dish with entomology pins through connective tissue and/or nerves to perform fine dissection.

We isolated the superior cervical ganglia (SCG) from mice by following a published protocol^50^. Briefly, mouse head was placed with the ventral side facing up. A midline incision to open the neck was performed, and skin and fat around trachea were cleared away. Parallel to the trachea, a pair of muscles were removed (both sides), and the carotid artery became visible. Once the “Y”-shaped bifurcation was identified in the carotid artery, the SCG was found and isolated using fine forceps.

### Ganglia dissociation and neuron isolation

Following SCSMG ganglia isolation from animals that were spleen-injected with Red Retrobeads, ganglionic neurons were dissociated. Briefly, in a 2 ml tube, isolated SCSMG or SCG were incubated in the dark in collagenase type I (2 mg/ml) in Leibovitz’s L-15 medium at 37°C, 120 rpm, for 30 min. Then, the ganglia were centrifuged at 400 g for 5 min. Supernatant was discarded and a new solation of collagenase type I (2 mg/ml) in 0.075% trypsin was added to resuspend the ganglia, which were incubated in the dark at 37°C, 120 rpm, for 30 min. Following centrifugation at 1000 g for 5 min at 4°C, ganglia cells were resuspended in 500 µl of Leibovitz’s L-15 medium (with DNAse 0.1 mg/ml). To assist the chemical dissociation (collagenase and trypsin), ganglia were gently pipetted up and down (physical dissociation) no more than 3 times throughout this protocol. After dissociation, an inverted fluorescence phase contrast microscope (Olympus IX71) was used to identify and collect individual labelled and unlabelled neurons for scRNAseq. Each neuron was collected by using a glass micropipette that was produced from capillary tubes (Borosilicate Glass from *SDR Scientific*; item #B150-86-15). A micropipette puller (Flaming/Brown micropipette Puller, model P-87) was used to make the capillaries, generating a tip around 40 µm. Looking under the microscope, and after positioning the glass micropipette on a dissociated single neuron, negative pressure by using the mouth of the experimenter was applied via silicone tubing connected to the micropipette to collect the neuron. Collected neurons were released in a 200 µl tube by applying a gentle positive pressure in the connected silicone tubing. The samples were snap frozen in a low temperature freezer (-80°C).

### Smart-seq analysis of isolated neurons

Single neurons were amplified using SMART-Seq HT Kit, as per manufacturer instructions, and quality control was performed using TapeStation electropherograms. Library preparation was performed using Nextera XT DNA library preparation kit as per manufacturer instructions. Sequencing was performed using an Illumina NextSeq 2000 P2 flow cell, 100 cycle run to an average depth of 4,836,605 reads per cell.

Whole transcriptome FASTQ files were aligned to either the mouse (GRCm39) or rat (mRatBN7.2) genome using STAR (version 2.7.9a)^51^. STAR was run with the –quantMode argument set to ‘TranscriptomeSAM’ to produce a transcriptome-co-ordinate BAM file. The output BAM files were split by read group to produce a separate BAM file for each sample. RSEM (version 1.16.1)^52^ was run to quantify transcript isoform expression and rsem-generate-data-matrix was run on the resulting files to produce merged expected counts matrices for mouse and rat samples.

For rat neurons, cells with less than 10% mitochondrial genes were retained for further analysis, resulting in 16 RFP^+^ and 11 RFP^-^ cells and 34034 detected genes. For mouse neurons, cells with less than 3% mitochondrial genes and cells filtered by expression of *Ptprc* (<0.5) and *Cd3g* (<0.1) were retained for further analysis, resulting in 56 RFP^+^ and 27 RFP^-^ SCSMG cells from naïve mice, 35 RFP^-^ SCG cells from naïve mice, and 3 RFP^+^ SCSMG cells from LCMV infected mice, with 40879 detected genes.

HUGO Gene Nomenclature Committee (HGNC) gene groups and gene ontology (GO) annotations were used to identify neuropeptide and neurotransmitter genes, and genes associated with synapses, nervous system development, ion channels and G protein coupled receptors. Genes distinguishing SCSMG and SCG neurons were determine in Seurat using FindMarkers() and data was visualised using the Seurat functions DoHeatmap() and VlnPlot(), and volcano plots generated with EnhancedVolcano (version 1.24.0).

### Interactome analysis

Information on interacting ligand-receptor pairs was obtained from the CellChatDB database^53^. Ligand-receptor interactions were considered to occur between spleen-innervating neurons and other cell types in the spleen (stromal or immune cell types) if the following conditions were met: (1) Genes encoding ligands or receptors in neurons and the corresponding partner receptors or ligands were expressed in at least 50% of spleen-innervating neurons with a z-score above 0.5.

### Single nucleus RNA sequencing of ganglia

Following SCSMG ganglia isolation from 5 naïve rats, 20 naïve mice and 20 LCMV-infected mice (day 8), nuclei were isolated from the frozen ganglia tissue using the Chromium Nuclei Isolation Kit (10X Genomics). Isolated nuclei were examined under a microscope and counted with a haemocytometer before loading. We counted a total of 14,000 nuclei from the rat isolated ganglia pool; 6,500 nuclei from the naïve mouse isolated ganglia pool; and 16,000 nuclei from the LCMV-infected mouse isolated ganglia pool. Each sample was loaded separately in individual lanes on the Chromium controller, for generation of gel bead-in-emulsions. Gene expression libraries were prepared using Chromium Next GEM Single Cell 3ʹ Kit v3.1 (10X Genomics), according to the manufacturer instructions. Library QC was performed using Tape station and Qubit. Sequencing was performed using an MGI DNBSEQ-G400 FCL, PE100.

### snRNA-seq analysis

Single nuclei sequencing samples were preprocessed and counted using the Cellranger(v7.0.0) count function against the mm10-2020-A (mouse) and MRatBN7.2 (rat) reference transcriptomes. Following this, each raw count matrix was further processed through CellBender^54^ (version 0.3.0) with the following flags: false positive rate (fpr) = 0.3 and epochs = 150 to remove ambient RNA contamination. The subsequent cleaned count matrices were further processed via R (version 4.4.1) and Seurat (version 5.1.0) using The University of Melbourne’s high performance computing system Spartan. Cells with at least 200 genes and less than 15% mitochondrial genes were retained for further analysis. Mouse SCG snRNA-seq data from Ziegler et al^25^ was downloaded from GEO (GSE231765) as a processed Seurat object (supplementary file *GSE231765_snSeq_integrated_res025_alra.RData.gz*). The snRNA-seq datasets (mouse and rat SCSMG and mouse SCG) were normalized using the SCTransform() function in Seurat. Samples were integrated using the IntegrateLayers() function with the Harmony integration method (version 1.2.1). Neuron clusters with high ambient expression of the long non-coding RNAs *Meg3, Fgf14 and Snhg11* were removed. Dimensionality reduction was performed by PCA and UMAP with 30 dimensions and cell clusters identified using the clustree package (version 0.5.1) and the Seurat functions FindNeighbors() with 30 dimensions and k.param = 30, and FindClusters() at a resolution of 0.4. Differentially expressed genes per cluster were identified using FindAllMarkers() or FindMarkers().

Neurons from mouse SCSMG and SCG, and rat SCSMG snRNA-seq were selected using the Subset() function before merging with mouse and rat neurons profiled by Smart-seq using merge(), SCTransform(), RunPCA(), IntegrateLayers(), FindNeighbors() and RunUMAP() with 40 dimensions and FindClusters() at a resolution of 0.2. Data was visualized using the Seurat fuctions VlnPlot(), FeaturePlot() and DimPlot() and dittoBarPlot() using dittoSeq (version 1.18.0).

Gene modules were added using AddModuleScore(), including the modules response_to_virus (*Ifitm3, Bst2, Adar, Ifit1, Ifit3, Ifi27l2a, Oasl2*) and antigen_presentation (*H2-Q7, H2-K1, H2-Q4, Tap2, Tap1, B2m, H2-D1, Tapbp*).

### scRNA-seq and Smart-seq integration

Raw datasets from Wei et al^26^ were downloaded from the Genome Sequence Archive (GSA: CRA030782; https://ngdc.cncb.ac.cn/gsa). 10x Chromium FASTQ files were processed and counted using Cellranger (v7.0.0) against a mm10-2020-A (mouse) reference using default parameters, which was modified by adding sequence for tdTomato, Cre, and FlpO to detect their transcripts. To remove ambient RNA contamination, raw count matrices were processed using CellBender^54^ (v0.3.0) remove-background with the following parameters: false positive rate (fpr) = 0.01, epochs = 150, learning rate = 1×10⁻⁶, expected number of cells = 35,000, total droplets included = 60,000, posterior batch size = 512, and empty droplet training fraction = 0.5. These parameters were selected to ensure stable convergence of the evidence lower bound (ELBO).

Denoised count matrices were imported into R (v4.0.3) and used for preliminary clustering with Seurat (v5.4.0). To standardise datasets annotated with different mouse genome assembly versions, the transcriptome was limited to the common set of 24,772 genes shared across the GRCm39 and mm10-2020-A. Major cell lineages were annotated based on canonical marker genes: *Scn7a* and *Fabp7* for satellite glia; *Tubb3* and *Th* for sympathetic neurons; *Pdgfra* and *Dcn* for fibroblasts; *Ptprc* for immune cells; *Pecam1* and *Flt1* for endothelial cells; *Rgs5* and *Pdgfrb* for mural cells; *Ncmap* and *Mpz* for Schwann cells; *Tpsab1* and *Cpa3* for mast cells; and *Hba-a1* and *Hba-a2* for erythrocytes. One cluster co-expressing *Fabp7* and *Th* was excluded as a doublet, as was one cluster with low marker gene expression and high mitochondrial content. Within each annotated cell type, cells deviating more than 3 median absolute deviations (MADs) from the median nCount_RNA in either direction were excluded as outliers. Cells with less than 15% mitochondrial reads were retained, and genes expressed in fewer than 3 cells were excluded. Putative doublets were identified independently within each sample using the scDblFinder R package (v1.20.2)^3^, with the clusters argument set to the preliminary lineage annotations and the samples argument set to sample ID.

Sympathetic neurons were extracted for downstream analysis. Log-normalized counts were calculated using the deconvolution strategy implemented by the computeSumFactors function in the scran R package (v1.34.0)^55^, followed by rescaled normalisation using the multiBatchNorm function in the batchelor R package (v1.22.0)^56^ to ensure comparable size factors across batches. The resulting log-transformed, rescaled expression values were used for subsequent integration and differential gene expression analysis. The top 2,000 highly variable genes (HVGs) were identified using FindVariableFeatures(), after excluding the sex-linked genes *Xist* and *Ddx3y* to avoid sex-driven clustering artifacts. Mitochondrial read percentage, a curated list of neuron-contamination genes, and cell cycle phase were regressed out using ScaleData(), and HVGs were centred and scaled across all cells.

The scRNA-seq and Smart-seq datasets were integrated using IntegrateLayers() with the Harmony integration method (v1.2.4)^57^. 12 principal components, selected based on an elbow plot, were used as input to FindNeighbors(), and cells were visualized using UMAP. Unsupervised clustering was performed with FindClusters() at a resolution of 0.1. Differentially expressed genes for each cluster were identified using FindAllMarkers().

### CRISPR–Cas9 editing of CD8^+^ T cells

CRISPR–Cas9 editing of naive CD8^+^ T cells was performed as previously described^58^. Single-guide RNAs (sgRNAs) targeting: Cxcr3 (5′-GAACAUCGGCUACAGCCAGG-3′, 5′-UGAGGGCUACACGUACCCGG-3′), Cxcr6 (5′-UCUGUACGAUGGGCACUACG-3′, 5′-UGUGCCAAAGACCCACUCAU-3′) and Cd19 (5′-AAUGUCUCAGACCAUAUGGG-3′, 5′-GAGAAGCUGGCUUGGUAUCG-3′) were purchased from Synthego (CRISPRevolution sgRNA EZ Kit). sgRNA/Cas9 ribonucleoprotein (RNP) complexes were generated by incubating 0.3 nmol of each sgRNA with 0.6 μl Alt-R S.p. Cas9 nuclease V3 (10 mg/mL; Integrated DNA Technologies, cat#1081059) for 10 min at room temperature. Naive CD8^+^ T cells were negatively enriched from the spleens of P14 mice by incubating cell suspensions with anti-CD4 (GK1.5), anti-CD11b (M1/70) anti-F4/80 (BM8), anti-Ter119 (TER-119) and anti-I-A/I-E (M5/114.15.2) monoclonal antibodies, followed by incubation with goat anti-rat IgG-coupled magnetic beads (Qiagen, cat#310107). Bead-bound cells were subsequently removed. A total of 1 × 10^7^ enriched T cells were resuspended in 20 μl of P3 (P3 Primary Cell 4D-Nucleofector X Kit; Lonza, cat#V4XP-3032), mixed with sgRNA/Cas9 RNPs and electroporated using a Lonza 4D-Nucleofector system (DN100). Cells were rested for 30 min in a 96-well plate before being directly transferred into recipient mice via intravenous injections (co-transfer of 1:1 ratio of sgCd19:Cxcr3 or sgCD19:Cxcr6 CRISPR-edited naive P14 cells, total of 5 × 10^4^ cells per mouse).

### Flow cytometry

In some experiments, mice were injected with 4 μg of anti-CD45 (clone 30-F11) antibody intravenously 3 min before euthanasia. Spleens were harvested from mice and placed into RPMI kept on ice. Cell suspensions were prepared by straining the organs through a steel mesh then filtered through 75 μm nylon mesh. Red blood cell lysis was then performed for spleens by resuspending the cell pellet in 2 ml of RBC Lysis buffer for 5 min, washed then resuspended in media. To perform flow cytometry on ganglia, the dissected tissue was digested in RP-2 buffer (collagenase D 2 mg/ml; DNAse 0.1 mg/ml; 0.8 mg/ml) at 37°C for 20 min. After digestion, samples were pipetted up and down to facilitate digestion. After centrifugation, cells were incubated with the antibody mix at 4°C for 30 min. Cells were washed and resuspended and Live/dead fixable Near-IR Dead Cell Stain Kit (Molecular Probes) was used to stain dead cells. For cell enumeration, Sphero calibration particles (BD) were added to samples before acquisition on either a Fortessa flow cytometer (BD) or Cytek® Aurora. Data were analyzed with FlowJo v10 software (TreeStar).

For flow cytometry of neurons, SCSMG from TH-cre.tdTomato mice were collected in ice-cold artificial cerebrospinal fluid (aCSF) supplemented with 10% fetal calf serum (FCS). Ganglia were digested in 100μl of pre-heated (37°C) digestion mixture. In brief, 1ml of digestion solution contains 140μl of TrypLE Express (Life Technologies), 200μl of papain (Worthington, LS003126; 37 mg/ml), 10μl of Collagenase D (Roche, 11088858001; 100 mg/ml), 20μl of Dispase II (Roche, 04942078001; 40mg/ml) and 20μl of DNAse I (Sigma-Aldrich, DN25; 5mg/ml) in aCSF. Ganglia were incubated for 15 min at 37°C, after which 1ml of aCSF+10%FCS was added, and samples were placed on ice. SCSMG were then mechanically dissociated with three needles of decreasing diameter (18G, 22G and 26G). The resulting suspension was then filtered through a 70 μm mesh cell strainer (Miltenyi Biotec). Cells were centrifuged at 1,600g for 5min, 4°C and stained. Cells were washed and analysed on a Cytek® Aurora flow cytometer. Data were analysed with FlowJo v10 software (TreeStar).

### Spleen AAV injections

TH-cre and wild-type mice were given 0.05mg/ml of analgesic Carprofen (Rimadyl 50mg/ml) supplied in their drinking water, starting the day prior to surgery and up to 48 h post-surgery. Mice were anaesthetised using isoflurane vapour (2.5-3% for induction and 1.5-2% used for maintenance), mixed with a 80:20 ratio of O2 and air. A fine glass capillary was pulled using a Narishige PC-100 Micropipette puller, backfilled with mineral oil and inserted into a nanoinjector attached to a control unit set (RWD, R-480 nanoliter microinjection pump). After clipping the fur of the left abdomen region of the mouse and sanitising the area with betadine/80% ethanol, mice were placed in a right lateral recumbent position on a surgical board for exposing the spleen. A 1cm incision of the skin and subsequently the peritoneum was made on the left flank area of the mouse, just below the rib cage. The spleen was exposed using sterilised forceps and a total of 5 2 μl injections were made across different locations in the spleen on a 45° angle using the nanoinjector. The spleens were infused with rAAV-hSyn-DIO-hM3D(Gq)-mCherry-WPRE-hGH polyA, AAV2/8 (2 x 10^12^ vg/mL BrainVTA) that was diluted 1:1 in PBS at a rate of 1000 nl/min. Following each infusion, the nanoinjector was kept in place for a retention time of 2-3 min to reduce risk of AAV backflow. The spleen was kept moist with sterile saline in between the injections. Following the injections, the incision was sutured (peritoneum and muscle first, and then the skin) and betadine was applied to the site. A subcutaneous injection of 0.05 mg/kg buprenorphine was given to mice immediately post-surgery as an analgesic and mice were placed on a heated pad until ambulatory.

### Acute LPS experiments and cytokine measurements

Lipopolysaccharide (LPS) from Escherichia coli O111:B4, purified by phenol extraction (Sigma-Aldrich L2630-10MGLPS) was injected i.p. at 10 mg/kg into mice 90 min before blood was collected by cardiac puncture. Blood was allowed to clot overnight, centrifuged 1000 g for 10 min at 4°C and serum collected. In experiments involving treatment of TH-DREADD mice or AAV targeted mice, 0.2 mg/kg Clozapine N-oxide (CNO) was injected i.p. 15-45 min prior to LPS treatment. Assessment of serum cytokines was performed using the LEGENDplex Mouse Anti-Virus Response Panel (13-plex) (BioLegend, Cat No. 740621) according to the manufacturer’s specifications. For assessment of spleen cytokines, spleens were collected and placed in a PBS solution containing 10 μl/ml protease inhibitor cocktail (P8340; Sigma-Aldrich, St. Louis, MS) and 1% BSA and then homogenised using a rotor-stator homogeniser. Spleen samples were centrifuged for 10 min at 10,000 g to collect the supernatant. ELISA was then performed on spleen supernatant and serum at appropriate dilutions for detection of TNF, IFNy and IL-10 using DuoSet ELISA development kits (R&D Systems: DY410, DY485, DY417) by following the manufacturer’s instructions.

### Intracellular cytokine staining

Following 90 min after LPS treatment, spleens were harvested and those allocated to flow cytometry were placed into a RPMI solution on ice. Samples were then digested with 2mg/ml collagenase in RP-2 media at 37°C for 30-35 min. Digested samples were broken up with the use of a pipette and then filtered through a 75μm nylon mesh. Following RBC lysis (5 min), suspensions were topped up with FACS buffer and filtered again through a 75μm nylon mesh, spun and resuspended in FACS buffer and plated in a 96-well ‘u-bottom’ plate. Following centrifugation, cells were resuspended in RP-10 solution containing Brefeldin-A (BD GolgiPlug 555029) and incubated for 4 h in a 36°C incubator. After washing with FACS buffer, samples were then stained with surface antibody cocktail and Zombie NIR fixable dead cell stain (Biolegend, 423106) at 4°C for 30 min. Washed samples were then permeabilised with Invitrogen fix/perm kit (eBioscience, 00-5123-43) for 40 min at 4°C, washed with permeabilization buffer and stained with intracellular cytokine antibody cocktail for 35-40 min at 4°C. Samples were washed and resuspended in FACS buffer containing calibration beads (BD, 556296). Samples were then acquired using the Cytek Aurora flow cytometer and analysed with FlowJo v10 software.

### Immunofluorescence and confocal microscopy

Mouse spleens were harvested and cut in halves or thirds and fixed in 4% paraformaldehyde (PFA) for 3-4 hrs at 4℃. Tissue was washed in PBS (3x5 min) before transferring to 30% Sucrose PBS-Azide for overnight cryoprotection at 4℃. Tissue was briefly submerged in 50:50 OCT:PBS-Sucrose/Azide before embedding in OCT (Sakura Finetek) and freezing at -30℃. Cryosections (12-50μm) were cut using a Leica cryostat (CM3050S) and air-dried at room temperature before proceeding to staining. A hydrophobic pap pen (Sigma; Z672548-1EA) was used around sections followed by application of Protein Block, Serum Free (Agilent Technologies; X090930-2) for 30 mins at room temperature. Protein block was removed and tissue was incubated in primary antibodies overnight at room temperature. The following day tissue was washed with 3x5 min PBS and incubated in secondary antibodies or fluorescently conjugated primaries for 1hr at RT. Tissue was then washed 3x5 min PBS. Coverslips (1, Knittel Glass, G417, 24 x 50 mm) were then mounted on sections using Prolong Diamond Antifade Mountant (Thermo Scientific; P36970). For nerve colocalization, 2 x 3x3 z-stack tile scans were imaged using either an LSM780 or LSM980 confocal microscope (Zeiss, Germany). The images were then processed with Imaris v10 (Oxford Instruments) and Fiji (ImageJ v1.5x). Co-localisation analysis and creation of a co-localisation channel for nerve quantitation was performed using Imaris, and the number of pixels in the co-localisation channel was enumerated using Fiji.

### Whole-mount ganglia clearing and staining

Isolated SCSMG from AAV targeted mice were post-fixed in 4% PFA for 3-4 h, washed and stored in PBS with 0.1% sodium azide. Whole ganglia were washed with PBS (3x 20min) and then incubated with blocking solution (PBS + 0.4% Triton X-100 + 10% normal donkey serum + 5% DMSO) for 4 hours on shaker at room temperature and left overnight at 4°C. Ganglia were incubated with primary antibody cocktail (Chicken anti-mCherry and Rabbit anti-cFOS), for 3 days in a 37°C orbital shaking incubator (70rpm). Following a set of washes (3x 30 min / overnight) with PBS + 0.3% Triton X-100, ganglia were incubated for 3 days in a 37°C orbital shaking incubator (70rpm) with secondary antibodies: goat anti-chicken AF594, and donkey anti-rabbit AF647). All antibodies were prepared in PBS + 0.1% sodium azide + 0.4% Triton X-100 + 3% normal donkey serum + 5% DMSO. Following another set of washes (3x 30 min / overnight) with PBS + 0.3% Triton X-100, ganglia were cleaned of contaminants, rinsed in PBS and dehydrated in a series of increased methanol gradients (25%, 50%, 75%, 2x 100%; 20 minutes each on shaker). Ganglia were delipidated in 100% dichloromethane (DCM) for 20-30 minutes, cleared with dibenzyl ether for 30 minutes and washed with ethyl cinnamate for mounting onto glass slides with coverslips (1, Knittel Glass, G417, 24 x 50 mm), which were then sealed with nail polish. Z-stack (2μm *z-*step) tile scans were imaged using the LSM980 confocal microscope (20x objective), and the images were then processed with Imaris v10 (Oxford Instruments).

### Myenteric plexus preparation and analysis

Jejunal tissue was dissected, opened longitudinally and pinned down for immersion-fixation in Zamboni’s fixative over night at 4°C. Tissue was washed with dimethyl-sulfoxide and PBS (3 x 10 min each) and myenteric plexus was dissected by removing the mucosal and submucosal layers as well as the circular muscle of the muscularis externa. Tissue was blocked using 10% normal horse serum in PBS containing 1% Triton-X 100 for 30 min at room temperature. After incubation with primary antibodies overnight at 4 °C, samples were washed 3 x 10 min with PBS before being incubated with secondary antibodies for 1 h at room temperature. After washing 3 x 10 min with PBS, tissue was mounted on glass slides using fluorescence mounting medium (Dako; S3023). Using a LSM980 confocal microscope (Zeiss, Germany), z-stack tile scans were acquired. A minimum of 1.4 mm^2^ per animal was analysed using ImageJ version 1.54f. TH signal was measured using an automatic local threshold and quantified in relation to total tissue area. Due to variable staining quality between experiments, values were normalised to non-infected controls of the respective experiment.

### Vibratome sectioning and clearing

Naïve or LCMV infected spleens were harvested and fixed in 2% PFA for 2-4 h at 4°C before being washed twice in PBS and embedded in 2% agarose. 150 µm tissue slices were obtained using a VT1200 S vibratome (Leica Biosystems) as previously described^59^. Each slice was placed into 2 ml SPW buffer overnight at 4°C with light agitation using a MR-1 Rocker-shaker (Fisher Biotec) before being transferred into washing buffer containing primary antibodies for 3 days at 4°C with light agitation. Slices were then washed for 2 h in washing buffer and another 2 times for 30 min in washing buffer. Washing buffer containing secondary antibodies was used to stain slices for a further 2 days at 4°C with light agitation. Slices were washed again as before and placed into FunGi clearing solution^60^ overnight at room temperature while agitated. A shallow well was created on glass slides using vacuum grease and slices mounted using FunGi clearing solution. Images were acquired imaged using a Zeiss LSM710 confocal microscope. Post-acquisition analysis was performed using Imaris.

### Sucrose-phosphate-glyoxylic acid (SPG) staining

20 μm sections were cut using a cryostat (Leica CM3050S) and placed immediately in SPG solution 1 containing malachite green for 3 seconds to create contrast then washed in SPG solution 2 for 3 seconds to wash out the excess malachite green. The sections were then dried using a hair dryer. Sections were covered in immersion oil and placed in an oven kept between 90°C-95°C. Excess oil was removed before covering with a glass coverslip and sealed with nail polish. Images were acquired using an SP8 fluorescence microscope (Leica), capturing between 440-520 nm and processed using Leica LAS X 1.9.

### Spleen tissue clearing and light sheet imaging

Whole spleen (cut in half to improve staining and clearing) was processed for tissue clearing using the Shanel protocol^11^ with some modifications. First, intracardiac perfusion was performed on anaesthetised mice (100 mg/kg ketamine; 2 mg/kg xylazine) using a syringe-infusion Pump 11 Elite (Havard Apparatus) at 2-3 ml/min speed. For lectin vessel labelling, anaesthetised mice were injected with 100 µl (1 mg/ml) of Tomato-Lectin (DyLight-649, L32472; Life Technologies/Thermo Fisher Scientific, Scoresby, Vic, Australia) into the left ventricle over a 20-30 second period and allowed to circulate for a further 2 min for uninfected mice or 3 min for LCMV infected mice, prior to perfusion. Mice were initially perfused with 10ml PBS followed by 15-20ml 4% paraformaldehyde (PFA) in PBS. Spleen was carefully removed and post-fixed in 4% PFA overnight followed by three washes in PBS and stored in PBS with 0.1% sodium azide. Following several washes in PBS, spleens were decolorised in 10% CHAPS/ 25% NMDEA (w/v) solution for 48 h at 37°C on an orbital shaker, with frequent changes of fresh solution. After incubation, spleens were washed in PBS (3x 20 min) and dehydrated in a series of increased ethanol gradients (50%, 70% and 100%; 2 h each on a tube rotator). Spleens were then incubated in a 2:1 dichloromethane (DCM)/ methanol solution overnight and then rehydrated with a series of 100%, 70%, 50% ethanol and distilled H_2_O (2 h each on a tube rotator). Spleens were incubated overnight in 0.5M acetic acid solution on a tube rotator (12 rpm), washed with dH_2_O twice, and incubated with Guanidine solution (4M guanidine hydrochloride, 0.05M sodium acetate and 2% Triton X100) for 6-8 h on rotation. Following another set of washes, spleens were blocked for 2 days with a DPBS solution containing 0.3% Triton X100, 10% DMSO and 10% normal donkey serum. Samples were then incubated with the following primary antibodies (TH 1:500 AB152 or CGRP 1:1000 C8198) for up to 6 days at 37°C on rotation at 70rpm. Following extensive washes in wash buffer (DPBS + 0.2% Tween-20 + 10 μg/ml heparin) up to 2 days, spleens were incubated in secondary antibody solution (donkey anti-rabbit IgG AF647: A32795) for 3 days at 37°C on rotation at 70rpm. All antibodies were prepared in a DPBS solution containing 0.3% Triton X100, 10μg/ml heparin, 3% DMSO, 10mM EDTA-Na pH8 and 5% normal donkey serum. Following another set of extensive washes in wash buffer, spleens were cleaned of contaminants and dehydrated (50%, 70%, 100% ethanol; 2 h each on tube rotator) and left overnight in 100% ethanol. Finally, samples were delipidated in 100% DCM (3x 1 hour) followed by incubation in 100% dibenzyl ether (3x 12 h) to match refractive index, on rotation at 12rpm. Spleens were transferred to ethyl cinnamate prior to imaging with the Ultramicroscope Blaze (LaVision Biotec, Miltenyi Biotec, Germany) using the 1.1x or 4x zoomable lens (MI PLAN, LaVision Biotec) with single or dual-sided illumination depending on the orientation. The numerical aperture was set at 0.16, exposure at 50-200 ms, sheet width at 100% and z-step at 2 μm. Raw acquired images were converted to imaris files (.ims) using the Imaris file converter (Bitplane) and all analyses was performed using Imaris software (Bitplane) including surface rending of nerve bundles and spleen surface using the *surfaces* function, and tracing of individual nerves using the *filament* function.

### Intracellular electrophysiological recordings

Animals were sacrificed by CO_2_ inhalation (SCG dissection) or by cervical dislocation (SCSMG dissection). SCSMG was dissected in mice as previously described^23^. Ganglia were removed and placed in ice-cold Krebs physiological saline, final molarities, with respect to ions (mM): 144 Na^+^, 4.76 K^+^, 1.2 Mg^2+^, 2.5 Ca^2+^, 128 Cl^−^, 25 equivalent HCO−3HCO3−, 1.06 H2PO4, H2PO4−, 1.2 SO2−4SO42−, and 11.1 glucose, bubbled with carbogen (95% O_2_-5% CO_2_.) Tissue was transported on ice to the electrophysiological setup and pinned onto a Sylgard-lined dish using Tungsten Pins (50-80 μm). Tissues were superfused with physiological Krebs solution at 32–34°C throughout experiment. Intracellular impalements into neurons were made using sharp glass microelectrodes, prepared from borosilicate glass capillaries (GC100F-15, 1 mm OD, 0.58 mm ID, Harvard Apparatus, Holliston, MA) using a P-97 Flaming/Brown Micropipette Puller (Sutter Instruments, California) and filled with 0.5 M KCl. A Leica micromanipulator was used to position the recording microelectrodes. Electrode impedances ranged from 70 to 200 MΩ. Data were recorded using an AxoClamp2B amplifier (Axon Instruments, California) in bridge mode using a 0.1 × LUT head-stage. Signals digitized using a PowerLab 4/26 (ADInstruments; PL2604) and analysed using Lab Chart Reader v8 (ADInstruments). Cells in ganglia that had a resting membrane potential (RMP) more hyperpolarized than −30 mV were used for analysis. RMP was determined in bridge balance mode before switching to direct current clamp (DCC) and changing V_holding_ to -60 mV. Current injections were made at 10pA increments until neurons fired an action potential (AP) and rheobase (pA) was recorded. To determine firing patten (tonic or phasic firing profile) neurons were injected with current (3-4x > threshold). Data is represented as normalised to non-infected mice with raw rheobase values in supplementary data.

### Stromal cell sorting

Isolation of stromal cells from mouse spleen was performed as previously described^28^. Briefly, spleens were harvested and placed into RPMI + 2.5% FCS solution on ice and then digested for 30 min at 37°C with an RP-2 buffer containing 1mg/ml collagenase, 0.8mg/ml dispase II and 0.1mg/ml DNAse I. A 27G needle/syringe was used to inject the solution across several locations of the spleen prior to incubation. Tissue was broken up with the use of a pipette and any remaining clumps were incubated again with digestion buffer at 37°C for a further 15 min. The digested solution was then filtered through a 75μm nylon mesh while topped up with ∼7-8ml of FACS buffer. Following RBC lysis (5 min), suspensions were topped up with FACS buffer and filtered again. After centrifugation, samples were resuspended in FACS buffer containing CD45 microbeads (Miltenyi Biotec; 1:10 for naïve and 1:5 for LCMV) and incubated at 4°C for 15 min. Samples were then passed through a LS column (Miltenyi Biotec) on a magnetic QuadroMACS^TM^ separator rack and the CD45 negative eluted portion was then collected after several washes of the column with FACS buffer. Eluted samples were stained with a stromal cell antibody cocktail at 4°C for 25 min, washed and resuspended in FACS buffer, and then sorted into pericyte and T cell zone FRCs populations using a Backman Coulter CytoFLEX SRT flow cytometer.

### Quantitative real time PCR

Total RNA was extracted from spleen or sorted stromal cells using RNeasy Plus Mini Kit (Qiagen) and converted to complementary DNA using the High-Capacity cDNA Reverse Transcription Kit (Thermo Fisher Scientific; Cat #4368813) according to the manufacturer’s instructions. Genes of interest were pre-amplified from cDNA using TaqMan PreAmp Master Mix (Thermo Fisher Scientific). Gene expression was analysed by real-time qPCR using Fast SYBR Green Master Mix (Thermo Fisher Scientific). Cycle-threshold values were determined for genes individually, and gene expression was normalized to the housekeeping genes Hprt and Gapdh (ΔCt) and presented as 2−ΔCt (arbitrary units). Primer sequences are as follows:

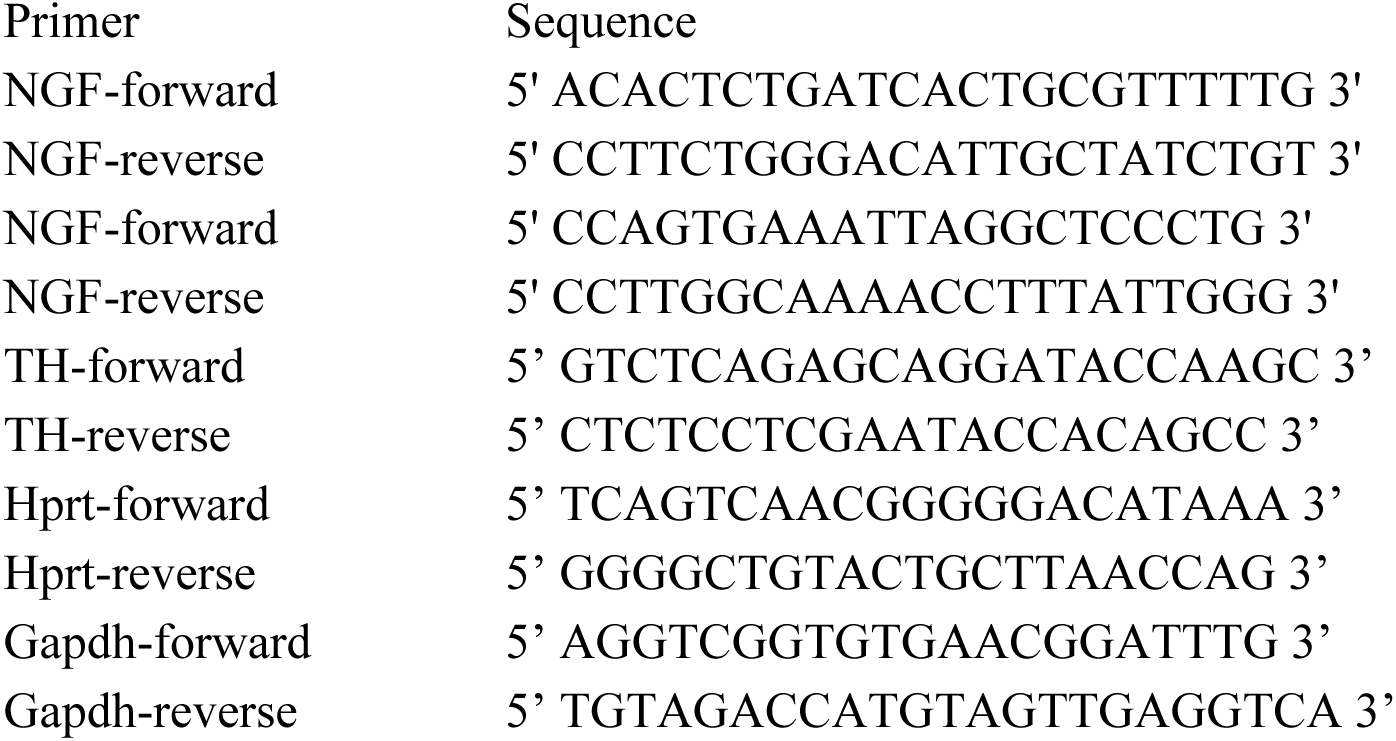

### Cell depletion

For NK cell depletion, mice were given 100 µg anti-NK1.1 antibody (PK136) i.p. two days prior to infection. For CD4 T cell depletion, mice were given 200 µg anti-CD4 antibody (GK1.5) i.p. 4 days and 1 day prior to infection. For CD8 T cell depletion, mice were given 200 µg anti-CD4 antibody (2.43) i.p. 4 days and 1 day prior to infection. Cell depletion was validated by flow cytometry in spleen. For TNF and IFNγ blockade, mice were injected with 200 μg anti-TNF (TN3-19.12) and 200 μg anti-IFNγ (XMG1.2) antibodies, or 200 μg isotype control antibodies (Armenian Hamster IgG clone PIP; Rat IgG2a Isotype clone 1-1) antibodies 4 h after LCMV infection, and again on days 3 and 6. For depletion of macrophages, mice were injected i.v. with 10 µl per g on days -1 and 1 of LCMV infection. Depletion was validated by flow cytometry of spleen.

### Noradrenaline ELISA

Spleens and plasma of LCMV infected and non-infected mice were collected. Spleens were weighed and homogenised with a Polytron homogeniser (Kinematica AG) for approximately 5-10 seconds (speed 19). Samples were assayed for noradrenaline using a Noradrenaline Research ELISA (LDN) as per manufacturer’s instructions. Data were acquired using a Multiskan Ascent microplate reader (Thermo Fisher Scientific).

### LCMV plaque assay

Confluent Vero cells were washed, trypsinised and diluted 1:20 and then plated in 6 well plates. The plates were incubated for 4 days until approximately 80% confluent. Plasma and spleens that were weighed were frozen and stored at -80°C. On the day of the assay, the samples were briefly thawed in a 37°C water bath and spleen homogenised with a Polytron homogeniser (Kinematica AG) for approximately 5-10 seconds (speed 19) then titrated in ten-fold dilutions. 200μL of the titrated samples was overlaid onto the confluent Vero cells and incubated for one hour. 1.5% agarose and 2X 199 media mixed in a 1:1 ratio was then added to each well and incubated for 4 days. A solution containing 1:1 1.5% agarose, 2X 199 media, and 2% Neutral Red solution (v/v) was overlaid onto each well. Plaques were enumerated using a dissecting microscope the following day.

### Statistical analysis

Graphs and statistics were generated using Prism 10 (GraphPad). Samples were tested for normality and two groups were compared using two-tailed Mann-Whitney test, unpaired two-tailed t test or Welch’s t-test if an F test indicated unequal variances. Multiple groups were analyzed with one-way ANOVA or Brown-Forsythe & Welch ANOVA for populations with unequal variances, or Kruskal-Wallis test with Dunn’s test for multiple comparisons, based on Gaussian distribution. All graphs depict means ± SEM. Details of statistical analysis are indicated in the figure legends and include the statistical test used. ns, non-significant, ∗p < 0.05, ∗∗p < 0.01, ∗∗∗ p < 0.001, ∗∗∗∗p < 0.0001.

## Data availability

Data from this study are included in the article and extended data. scRNA-seq data from mouse and rat neurons are available at the Gene Expression Omnibus under the accession number GSE283204. snRNA-seq data from mouse and rat SCSMG are available under the accession numbers GSM8657805, GSM8657806 and GSM8657807.

## List of antibodies used in this study

| Reagent or Resource | Clone | Source | Identifier | Dilution | Use |
| --- | --- | --- | --- | --- | --- |
| PE-Cy7 anti-mouse CD45.1 | A20 | Thermo Fisher | Cat# 25-0453-82;<br>RRID: AB_469629 | 1/200 | Flow/IF |
| Brilliant Violet 785™ anti-mouse Ly6C | HK1.4 | Biolegend | Cat# 128041, RRID:<br>AB_2565852 | 1/400 | flow |
| APC-Cy7 anti-mouse TCRbeta chain | H57-597 | BD Biosciences | Cat# 560656, RRID:<br>AB_1727574 | 1/200 | flow |
| APC anti-mouse CD64 (FcγRI) | X54-5/7.1 | Biolegend | Cat# 139306, RRID:<br>AB_11219391 | 1/100 | flow |
| AlexaFluor™700 anti-mouse MHC ClassII (I-A/I-E) | M5/114.15.2 | Thermo Fisher | Cat# 56-5321-82,<br>RRID: AB_494009 | 1/400 | flow |
| BDHorizon™BUV805 anti-Mouse CD4 | RM4-5 | BD Biosciences | Cat# 569193 | 1/400 | flow |
| BDOptiBuild™BV750 anti-Mouse CD8b | H35-17.2 | BD Biosciences | Cat# 747505, RRID:<br>AB_2872172 | 1/200 | flow |
| Spark Blue 550 anti-mouse Gr-1 | RB6-8C5 | Biolegend | Cat# 108468, RRID:<br>AB_2860618 | 1/200 | flow |
| AlexaFluor647 anti-mouse Ly6G | 1A8 | Biolegend | Cat# 127610, RRID:<br>AB_1134159 | 1/400 | flow |
| Brilliant Violet 661 anti-human/mouse CD11b | M1/70 | BD Biosciences | Cat# 612977, RRID:<br>AB_2870249 | 1/400 | flow |
| PE/Fire™ 810 anti-mouse CD11c | QA18A72 | Biolegend | Cat# 161106, RRID:<br>AB_2904307 | 1/400 | flow |
| APC/Cyanine7 anti-mouse CD19 | 1D3 | Biolegend | Cat# 115530; RRID:<br>AB_830707 | 1/200 | flow |
| Streptavidin PE-Cy7 |  | BD Biosciences | Cat# 557598; RRID:<br>AB_10049577 | 1/400 | flow |
| Pacific Blue™ anti-mouse H2-Kb | AF6-88.5 | Biolegend | cat#116514,<br>AB_1967129 | 1/200 | flow |
| PerCP anti-mouse CD45, clone | 30-F11 | BD Biosciences | cat#557235,<br>AB_396609 | 1/200 | flow |
| PerCP/Cyanine5.5 anti-mouse CD31 | 390 | Biolegend | cat#102420,<br>AB_10613644 | 1/200 | flow |
| Alexa Fluor 488 anti-mouse Podoplanin | 8.1.1 | eBioscience | cat#53538182,<br>AB_1106990 | 1/200 | flow |
| BUV496 anti-mouse CD45R/B220 | RA3-6B2 | BD Biosciences | Cat# 612950,<br>RRID:<br>AB_2870227 | 1/300 | Flow |
| BD OptiBuild™ BUV563 anti-mouse NK1.1 | PK136 | BD Biosciences | Cat# 741233,<br>RRID:<br>AB_2870785 | 1/200 | Flow |
| BV421 anti-mouse TNFa | MP6-XT22 | Biolegend | Cat# 506328,<br>RRID:<br>AB_2562902 | 1/200 | Flow |
| BV421 anti-mouse CD31 | 390 | Biolegend | Cat# 102424,<br>RRID:<br>AB_2650892 | 1/500 | Flow |
| eFlour™ 450 anti-mouse Ly6G | 1A8-Ly6G | eBioscience | Cat# 48-9668-82,<br>RRID: 2637124 | 1/400 | Flow |
| BV480 anti-mouse IFNγ | XMG1.2 | BD<br>Biosciences | Cat# 566097,<br>RRID:<br>AB_2739549 | 1/400 | Flow |
| BV570 anti-mouse Ly6C | HK1.4 | Biolegend | Cat#128030,<br>RRID:<br>AB_2562617 | 1/400 | Flow |
| BV605 anti-mouse CD11c | N418 | Biolegend | Cat# 117333,<br>RRID:<br>AB_11204262 | 1/200 | Flow |
| BV605 anti-mouse CD140a | APA5 | Biolegend | Cat# 135916,<br>RRID:<br>AB_2721548 | 1/200 | Flow |
| BD OptiBuild™ BV650 anti-mouse CD157 | BP-3 | BD<br>Biosciences | Cat# 740611,<br>RRID:<br>AB_2740311 | 1/500 | Flow |
| BV785 anti-mouse CD8a | 53-6.7 | Biolegend | Cat# 100750,<br>RRID:<br>AB_2562610 | 1/200 | Flow |
| PerCP-Cy5.5 anti-mouse CD21/CD35 | 7G6 | BD<br>Biosciences | Cat# 562797,<br>RRID:<br>AB_2737802 | 1/100 | Flow |
| PE anti-mouse CD3e | 500A2 | eBioscience | Cat# 12-0033-82, RRID:<br>AB_842786 | 1/400 | Flow |
| PE-Vio615 anti-mouse CD106 (VCAM-1) | REA971 | Miltenyi<br>Biotec | Cat# 130-116-223,<br>RRID:<br>AB_2727416 | 1/50 | Flow |
| PE-Cy7 anti-mouse Podoplanin | 8.1.1 | eBioscience | Cat#25-5381-82,<br>RRID:<br>AB_2573460 | 1/500 | Flow |
| AF647 anti-mouse F4/80 | BM8 | Biolegend | Cat# 123122, RRID: AB_893480 | 1/200 | Flow |
| BD Horizon™ R718 anti-mouse CD45 | 30-F11 | BD Biosciences | Cat# 567075, RRID: AB_2916421 | 1/100 | Flow |
| BD Horizon™ R718 anti-mouse TER119 | TER-119 | BD Biosciences | Cat# 567363, RRID: AB_3683801 | 1/100 | Flow |
| purified Rabbit Anti-S100 | Polyclonal | Agilent | IR50461-2 | 1/500 | IF |
| Purified Rabbit anti-tyrosine hydroxylase | Polyclonal | Millipore | Cat# AB152, RRID: AB_390204 | 1/500 | IF |
| Purified Chicken anti-Tyrosine Hydroxylase | Polyclonal | Abcam | Cat# ab76442, RRID: AB_1524535 | 1/800 | IF |
| Purified Rabbit anti-beta III Tubulin | Polyclonal | Abcam | Cat# AB18207, RRID: AB_444319 | 1/800 | IF |
| Purified Rabbit anti-synaptophysin | Polyclonal | Abcam | Cat# AB14692, RRID: AB_301417 | 1/500 | IF |
| Purified rabbit anti-CGRP | Polyclonal | Millipore | Cat# C8198, RRID: AB_259091 | 1/1000 | IF |
| Alexa Fluor 647 anti-mouse CD31 | WM59 | Biolegend | Cat# 102516, RRID: AB_2161029 | 1/200 | IF |
| Purified Chicken anti-mCherry | Polyclonal | Abcam | Cat# ab205402, RRID: AB_2722769 | 1/800 | IF |
| Purified Rabbit anti-cFOS | Polyclonal | Abcam | Cat# ab190289, RRID: AB_2737414 | 1/400 | IF |
| Pacific Blue™ anti-mouse/human CD45R/B220 | RA3-6B2 | BioLegend | Cat# 103227; RRID: AB_492876 | 1/200 | IF |
| APC anti-mouse CD146 | ME-9F1 | Biolegend | Cat# 134712; RRID: AB_2563088 | 1/200 | IF |
| AlexaFluor700 anti-mouse CD3 | 17A2 | Thermo Fisher | Cat# 56-0032-82, RRID: AB_529507 | 1/200 | IF |
| Rabbit-anti-CD3e | Polyclonal | Agilent | Cat# A0452, RRID: AB_2335677 | 1/200 | IF |
| Alexa Fluor 647 anti-mouse CD8a | 53-6.7 | Biolegend | Cat# 100724, RRID: AB_389326 | 1/100 | IF |
| Alexa Fluor 594 InVivoMab anti-LCMV nucleoprotein antibody | VL-4 | Bio X cell | Cat# BE0106; RRID: AB_10949017 | 1/200 | IF |
| Alexa Fluor 488 Goat anti-chicken | Polyclonal | Thermo Fisher | Cat# A11039, RRID: AB_2534096 | 1/800 | IF |
| Alexa Fluor 488 Donkey-anti-Chicken | Polyclonal | Jackson ImmunoResearch | Cat# 703-545-155, RRID: AB_2340375 | 1/800 | IF |
| Alexa Fluor 488 Donkey anti-rabbit | Polyclonal | Thermo Fisher | Cat# A21206, RRID: AB_2535792 | 1/500 | IF |
| Alexa Fluor 594 Donkey anti-rabbit | Polyclonal | Thermo Fisher | Cat# A32754, RRID: AB_2762827 | 1/800 | IF |
| Alexa Fluor 594 Donkey-anti-Human | Polyclonal | Jackson ImmunoResearch | Cat# 709-585-149, RRID: AB_2340572 | 1/800 | IF |
| Alexa Fluor 594 Goat anti-chicken | Polyclonal | Thermo Fisher | Cat# A32759, RRID: AB_2762829 | 1/500 | IF |
| Alexa Fluor plus 647 Donkey-anti-Rabbit | Polyclonal | Thermo Fisher | Cat# A32795, RRID: AB_2866496 | 1/800 | IF |

## Notes

### Competing Interest Statement

The authors have declared no competing interest.

